# Translational profiling uncovers a tonoplast sugar transporter essential for vascular system development and cell wall composition

**DOI:** 10.64898/2026.04.29.721592

**Authors:** Beate Hoffmann, Françoise Vilaine, Alexandra Launay-Avon, Deyan Markovic, Stephen Lima, Mohamad Yassine, Audrey Hulot, Nadia Bessoltane, Christine Paysant-Le Roux, Sylvie Dinant, Etienne Delannoy, Rozenn Le Hir

## Abstract

The inflorescence stem of *Arabidopsis thaliana* is a powerful model to study vascular development and carbon allocation, yet the translational landscape underlying xylem differentiation remains poorly defined. Here, we address this gap using tissue-specific Translating Ribosome Affinity Purification sequencing (TRAP-seq) to resolve the translatomes of xylem vessels, xylem parenchyma, and interfascicular fibers.

We show that the very early stage of metaxylem differentiation involves coordinated activation of primary metabolism, plastid functions, and cell wall biosynthesis, supporting the metabolic demands of secondary wall formation. Xylem parenchyma displays enrichment in hormone signaling, stress, and immune pathways, consistent with a role in integrating environmental and metabolic cues. Interfascicular fibers exhibit increased translation of ribosomal proteins and spliceosome components, pointing to high translational activity and a role for alternative splicing during late development and programmed cell death.

We further identify the tonoplast sugar transporter SUGAR WILL EVENTUALLY BE EXPORTED TRANSPORTER 2 (SWEET2) as a key regulator of stem radial growth. Preferentially translated in early stage of metaxylem differentiation, SWEET2 acts as a rate-limiting component of vacuolar sugar exchange, controlling cytosolic hexose availability for secondary wall biosynthesis. By contrast, SWEET16 and SWEET17 exert more specialized, tissue-specific functions. Together, our findings establish subcellular sugar partitioning as a central determinant of vascular development and highlight the power of translatome profiling.

## INTRODUCTION

The inflorescence stem of *Arabidopsis* is organized as a cylindrical structure in which concentric tissue layers ensure protection, transport, mechanical support, and environmental sensing. The outermost layer, the epidermis, consists of a single layer of cells covered by a cuticle that limits water loss, protects against pathogens, and contributes to overall mechanical integrity (Shi et al. 2021; Asaoka et al. 2023, 2024). Beneath it, the cortex is composed of several layers of parenchyma cells that participate in defining stem geometry and mechanical properties and can adapt to environmental cues such as mechanical stress or gravity (Paul-Victor and Rowe 2011; Shi et al. 2021; Shinohara et al. 2024). The innermost cortical layer differentiates into the starch sheath, a starch-rich endodermis-like tissue involved in gravity sensing and later serving as a major source of interfascicular cambium, thereby linking primary and secondary growth processes (Altamura et al. 2001; Sehr et al. 2010; Shi et al. 2021; Turley and Etchells 2022). Vascular bundles are composed of phloem on the outer side, xylem on the inner side, and a procambial or cambial zone in between that acts as a stem cell niche for vascular tissue production (Benítez and Hejátko 2013; De Rybel et al. 2016; Shi et al. 2021). The phloem, composed of sieve elements and companion cells, ensures the transport of sugars and signaling molecules, while the phloem cap becomes lignified and reinforces mechanical strength (Altamura et al. 2001; Shi et al. 2021). Xylem tissue consists of different cell types: (i) tracheary elements (including vessels and tracheids), (ii) xylary fibers and (iii) xylem parenchyma cells (Lehmann and Schneider 2025). Xylem vessels and fibers possess thick, lignified secondary walls that enable efficient water transport and provide strong axial support (Altamura et al. 2001; Zhong et al. 2001; Kaneda et al. 2011; Asaoka et al. 2024). Associated xylem parenchyma cells, which have received comparatively little attention relative to other xylem cell types, are present in and around the vascular bundles. Arabidopsis stems contain at least two distinct xylem parenchyma populations: thin-walled parenchyma cells present at the bottom of the vascular bundle (Altamura et al. 2001; Smith et al. 2017a; Hoffmann et al. 2020), and xylem parenchyma closely associated with vessels and fibers that develop thicker but relatively unlignified walls (Baghdady et al. 2006). They provide storage and metabolic support to conductive elements and contribute to transport functions and cell wall or lignin metabolism (Altamura et al. 2001; Kaneda et al. 2011; Smith et al. 2013, 2017a; Shi et al. 2021). Furthermore, Smith et al. (2017) proposed the “good neighbor hypothesis,” suggesting that xylem parenchyma cells actively mediate monolignol transport to developing vessels during lignification, highlighting their underappreciated contribution to vessel development. Between vascular bundles, the interfascicular region initially consists of parenchyma cells, but at later stages of floral stem development, several inner layers differentiate into interfascicular fibers with thick lignified walls, forming a nearly continuous supportive ring essential for maintaining upright growth (Altamura et al. 2001; Zhong et al. 2001; Little et al. 2002; Paul-Victor and Rowe 2011; Asaoka et al. 2024). As secondary growth progresses, outer interfascicular cells also lignify, and an interfascicular cambium can develop, often derived from starch sheath cells, contributing to the formation of secondary xylem and phloem and further strengthening the stem (Altamura et al. 2001; Little et al. 2002; Sehr et al. 2010; Turley and Etchells 2022). Finally, at the center of the stem, the pith consists of parenchyma cells that play key roles in storage and structural organization, influencing both stiffness and flexibility of the stem (Paul-Victor and Rowe 2011; Shi et al. 2021; Asaoka et al. 2024; Shinohara et al. 2024)

Decades of research in both woody and herbaceous species have elucidated many regulatory networks, including identification of master genes/transcription factors governing xylem differentiation and development, particularly the formation of secondary cell walls (SCWs) and, to a lesser extent, the development of interfascicular fibers (reviewed in Hunziker and Greb 2024; Lehmann and Schneider 2025). Beyond transcription factors, studies have highlighted the critical role of primary metabolism in xylem development and SCW synthesis. For example, Ohtani et al. (2016) demonstrated that protoxylem vessel element differentiation is accompanied by metabolic reprogramming, likely facilitating the production of polysaccharides and lignin for SCW construction. Furthermore, the SUGAR WILL EVENTUALLY BE EXPORTED TRANSPORTER 11 (SWEET11) and SWEET12 transporters have been shown to be essential for proper xylem development and SCW formation in Arabidopsis stems (Le Hir et al. 2015; Dinant et al. 2019). Pinard et al. (2019) also revealed in *Eucalyptus* that the exchanges of primary metabolites between subcellular compartments, plastids and mitochondria, is tightly coordinated during xylogenesis. Subsequent work in Arabidopsis and *Populus tomentosa* further established that vacuole-to-cytosol sugar transport, mediated by SWEET family members, is also required for both xylem development and SCW composition (Aubry et al. 2022; Lu et al. 2025). These studies also underscore the importance of xylem parenchyma cells in these processes.

Despite these studies, the molecular mechanisms linking sugar metabolism and transport to the functions of the different xylem cell types and interfascicular fibers remain poorly understood. While tissue-specific and single-cell transcriptomic analyses of the *Arabidopsis* inflorescence stem have provided valuable insights (Shi et al. 2021; Lee et al. 2025), transcriptomic analyses fail to account for translational regulation, which plays an important role in gene expression control, particularly for genes involved in sugar metabolism or regulated by sugars (Gamm et al. 2014). To address these gaps, we employed the Translating Ribosome Affinity Purification (TRAP) assay using three cell-type-specific promoters: *LACCASE 17/LAC17* (driving expression in interfascicular fibers) (Berthet et al. 2011), *VASCULAR-RELATED NAC-DOMAIN 6*/*VND6* (driving expression in xylem vessels during metaxylem differentiation) (Kubo et al. 2005; Zhong et al. 2008), and *SWEET17* (driving expression in xylem vessels during metaxylem differentiation, xylary fibers and xylem parenchyma cells) (Aubry et al. 2022; Hoffmann et al. 2022). This approach enabled us to generate a high-resolution, cell-specific translatome, offering new insights into the molecular actors driving xylem development and SCW synthesis. Furthermore, we identified SWEET2, a tonoplast sugar transporter as a new molecular actor involved SCW formation in both xylem tissues and interfascicular fibers.

## RESULTS

Previous work has been deciphering the tissular fingerprint of several vascular cell types in Arabidopsis inflorescence stem transcriptomic analysis (Shi et al. 2021). However, translatomes can provide additional information regarding translational regulations, for example for gene’s transcripts that are associated with ribosomes as a result of repetitive translational reinitiation (Mustroph et al. 2009). In order to gain more insight into the translational machinery taking place in different xylem cell types and in interfascicular fibers, we produced Arabidopsis lines expressing the tagged Ribosomal Protein L18 (RPL18) under the expression of different promoters which have been described previously to display tissular specificity. We selected the promoter of *LAC17*, initially described as specific to interfascicular fibers (Berthet et al. 2011). Nonetheless, subsequent studies showed that *LAC17* is also expressed throughout protoxylem tracheary elements differentiation (a term encompassing both tracheids and xylem vessels) (Schuetz et al. 2014). This was further supported by work from Blaschek et al. (2023), who demonstrated *LAC17* expression during tracheary element differentiation. Moreover, analysis of multiple *laccase* mutant lines indicated that although lignin composition is more strongly affected in interfascicular fibers, *LAC17* also contributes to lignification in xylary fibers (Blaschek et al. 2023). Together, these findings suggest that *LAC17* is expressed and acts mostly in interfascicular fibers but also in tracheary elements and xylary fibers. In addition, we used the promoter of *VND6*, which drives expression specifically to xylem vessels undergoing metaxylem differentiation (Kubo et al. 2005; Zhong et al. 2008), and the promoter of *SWEET17*, reported to drive expression in xylem cells including xylem vessels (at the metaxylem stage), xylary and interfascicular fibers, and xylem parenchyma cells (Aubry et al., 2022). Based on their known expression patterns, we therefore expect that the comparison between them will yield to novel information regarding the role of xylem vessels (at the metaxylem stage), xylem parenchyma cells and interfascicular fibers in Arabidopsis inflorescence stems.

### Vascular bundle cell type-specific translational profiles can be determined in inflorescence stem

To verify the tissue specificity of the translatomes, we checked the activities of the different promoter used by generating transgenic Arabidopsis lines expressing a fusion protein between *RPL18* and the gene reporter *GUS* under the control of each promoter (*pGENE:HF-RPL18-GUS*). At the bottom part of the stem, a blue signal, corresponding to the expression domain of *pLAC17*, was observed mostly in interfascicular fibers as previously shown in Berthet et al. (2011) (Fig. 1A-B). In addition, some signal could also to observed in differentiating metaxylem vessel elements (Fig. 1B, black arrowheads). Regarding the expression pattern observed in *pVND6:HF-RPL18-GUS*, a blue signal was mainly observed in xylem vessels undergoing metaxylem differentiation close to the cambium layer (Fig. 1C-D). A GUS activity was detected in xylem parenchyma cells and differentiating metaxylem vessels in *pSWEET17:HF-RPL18-GUS* line (Fig. 1E), confirming the expression pattern previously reported for the *pSWEET17:GUS* line, both constructs sharing the same promoter length (Aubry et al. 2022). Considering our earlier observation of differential SWEET17 expression using a translational GUS fusion line with a longer promoter region, which varied depending on the vascular bundle type (Hoffmann et al. 2022), we also examined the *SWEET17_pro_:HF-RPL18-GUS* in the L-type vascular bundle. In this case, expression was specifically restricted to xylem parenchyma cells characterized by unlignified, thickened cell walls (Fig. 1F). These observations reveal overlapping GUS activiy of *pLAC17:HF-RPL18-GUS*, *pSWEET17:HF-RPL18-GUS* and *pVND6:HF-RPL18-GUS* in differentiating metaxylem vessels while also highlighting tissue-specific expression patterns. Specifically, *pSWEET17:HF-RPL18-GUS* is the only line expressed in xylem parenchyma cells, whereas *pLAC17:HF-RPL18-GUS* is uniquely expressed in interfascicular fibers.

**Figure 1.**
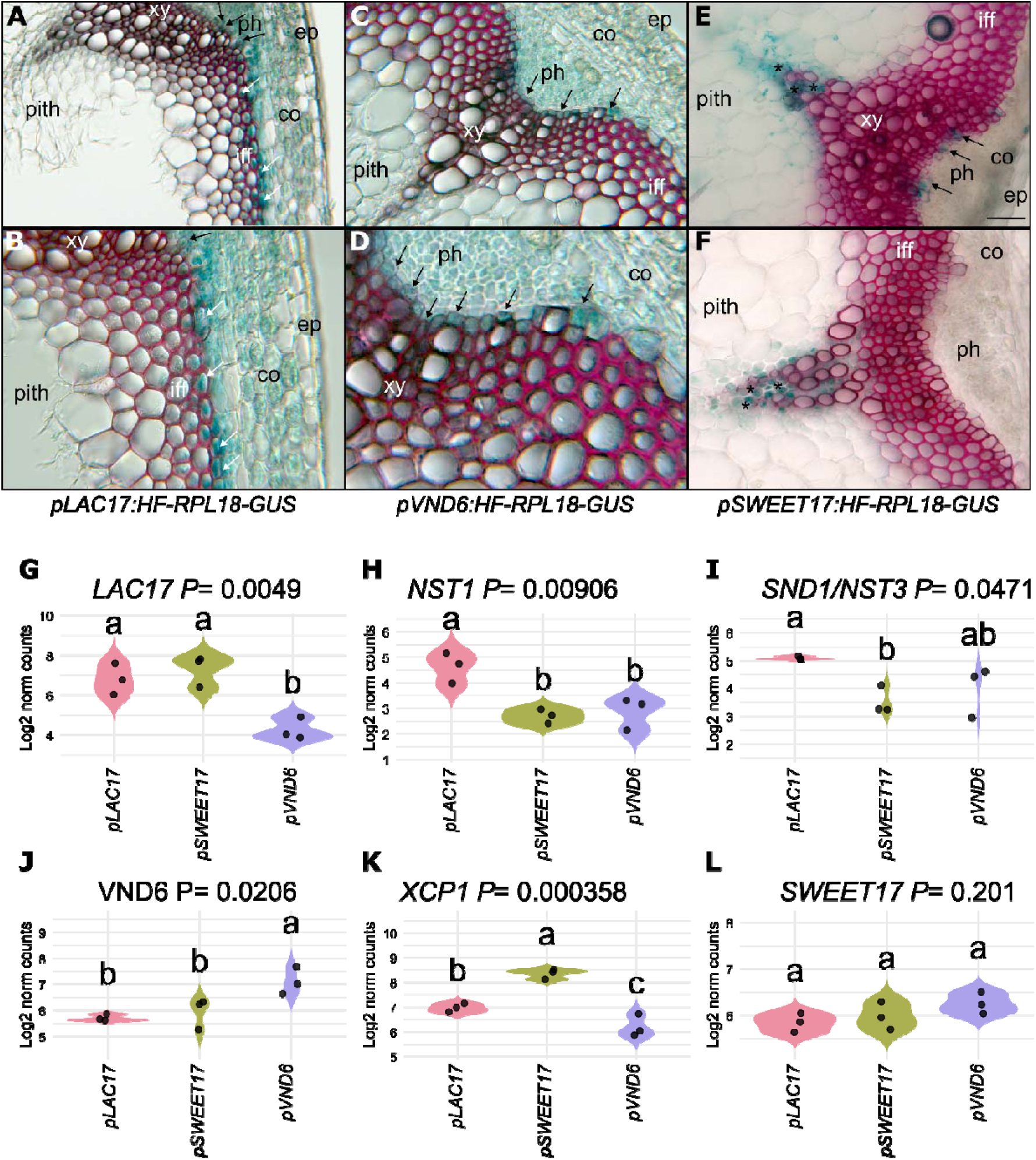
Expression patterns of *HF-RPL18-GUS* in the Arabidopsis stem. (A-F) Transversal sections taken at the bottom part of the stem in a region where elongation growth has finished but where thickening of the secondary cell wall was still ongoing of *pLAC17:HF-RPL18-GUS* line (A-B), *pVND6:HF-RPL18-GUS* (C-D) and *pSWEET17:HF-RPL18-GUS* (E-F). (B and D) close-up of panels (A) and (C), respectively. (E-F) Expression of *pSWEET17:HF-RPL18-GUS* in M-type (E) or L-type vascular bundle. Black arrows points to cells showing blue GUS staining in differentiating metaxylem vessel elements, asterisks indicate xylary parenchyma cells and white arrows points to differentiating interfascicular fibers. Lignin is colored pink after phloroglucinol staining. The intensity of the pink color indicated the level of lignification of the xylem tissue. co: cortex; ph: phloem, xy: xylem. Scale bar = 50 µm. (G-L) Violin plots showing log_2_-transformed normalized gene reads counts of the *LAC17* (G), *NST1* (H), *SND1/NST3* (I), *VND6* (J), *XCP1* (K) and *SWEET17* (L) among the different translatomes. Each black dot corresponds to a biological replicate (n = 3). A one-way ANOVA combined with Tukey’s comparison post-test has been made to compare the different translatomes. The different letters indicate significant differences. The *P*-value for the comparison between the different translatomes is indicated on the graph.

Following immunopurification of the *pGENE:HF-RPL18* lines, translatomes associated with the different promoters were generated using Illumina sequencing. Among the 37,051 Arabidopsis protein-coding and noncoding genes (Cheng et al., 2017), approximately 40% of transcripts were found to be associated with ribosomes—and thus actively translated—in each of the analyzed cell types (Supplementary Table S1). These proportions confirmed comparable translational activity across all cell types (Supplementary Table S1). Then, principal component analysis (PCA) was applied to the nine RNA-seq datasets obtained (Supplementary Fig. S1 and Supplementary Data Set 1). We observed a clear separation of the three tissues types (*pLAC17*-associated, *pSWEET17*-associated and *pVND6*-associated expression domains) within the projection of the two first principal component planes which gathered 42% of the total variance (Supplementary Fig. S1). Especially the first component clearly separates the *pVND6*-associated translatome from the two others and the second component separates *pSWEET17*-associated translatome from the *pLAC17*-associated translatome (Supplementary Fig. S1).

Further, we examined the expression levels of genes corresponding to the promoters used to drive expression of the HF-RPL18 fusion protein (Fig. 1G-L). As expected, *LAC17* expression was enriched in the *pLAC17*-associated translatome compared to the *pVND6*-associated translatome (Fig. 1G). Nonetheless, similar *LAC17* expression level was observed in the *pSWEET17*-associated translatome compared to the *pLAC17*-associated translatome (Fig. 1G). Importantly, the *NAC SECONDARY WALL THICKENING PROMOTING FACTOR 1* (*NST1*) and *SECONDARY WALL-ASSOCIATED NAC DOMAIN 1* (*SND1/NST3*) genes, well-established markers of fiber cells in Arabidopsis stems (Schuetz et al. 2012; Shi et al. 2021), were significantly enriched in the *pLAC17*-associated translatome (Fig. 1H-I), likely supporting that this dataset represents the translational profile of fiber cells. Conversely, *VND6* expression was highly enriched in the *pVND6*-associated translatome relative to the *pLAC17*- and *pSWEET17*-associated ones (Fig. 1J). Surprisingly, the expression of *XCP1*, a marker gene for xylem vessel differentiation (Funk et al. 2002), was significantly lower in *pVND6*-associated translatomes while it was enriched in the *pSWEET17*-associated translatome and, to a lesser extent, in *pLAC17*-associated translatomes (Fig. 1K). Finally, *SWEET17* expression was relatively uniform across all translatomes (Fig. 1L). The high levels of *SWEET17* in *pVND6*-associated translatomes is consistent with previously reported expression patterns in developing xylem vessels using transcriptional GUS reporter (Aubry et al., 2022). Taken together, these results suggest that the *pSWEET17*-associated translatome represents a composite of developing xylem vessels and xylem parenchyma cells. Furthermore, the detection of *SWEET17* expression in *pLAC17*-associated cell types points to a potential role for this transporter in interfascicular fibers development or function, consistent with previous report showing SWEET17 expression in interfascicular fibers (Hoffmann et al. 2022). The apparent discrepancy between the absence of *SWEET17* promoter-driven signal in interfascicular fibers and its detection in the translatome could be explained by the existence of regulatory elements or post-transcriptional control that are absent in the transcriptional GUS construct, while the translational fusion captures the full expression pattern at the protein level. As a result, *SWEET17* expression in interfascicular fibers may be captured in the translatome but not in the analyzed GUS sections (Aubry et al. 2022). Thus, despite the clear separation of the translatomes (Supplementary Fig. 1), a certain level of overlap exists between the different datasets as illustrated by the promoter expression domains and the expression of some marker genes (Fig. 1).

### Pairwise comparisons of translatome profiles reveal key enrichments in distinct functional categories

Given the partial overlap and ambiguity in gene assignment, we next performed a more detailed differential expression analysis and GO functional enrichment analyses for each pairwise comparisons, to better resolve the biological significance of these features. A differential expression analysis was performed using the DiCoExpress package in R (Lambert et al. 2020; Baudry et al. 2022) and allowed to identify a total of 1289 genes differentially expressed when comparing *pLAC17*- and *pVND6*-associated translatome (Supplementary Fig. S2A and Supplementary Data Set 2). The comparison between *pLAC17*- and *pSWEET17*-associated translatomes yielded to 858 DEGs and those between *pSWEET17*- and *pVND6*-associated translatomes to 748 DEGs (Supplementary Fig. S2A and Supplementary Data Set 2). To further analyze these datasets, we used the in-house “Arabidopsis Gene Network Explorer”- *ARAGNE* R app (https://forge.inrae.fr/aragne/ARAGNE), an all-in-one shiny interface for fast data exploration, user friendly GO and KEGG enrichment analysis and easy Cytoscape-based representation of gene regulatory network (GRN) (see ‘Material and Methods’ section for more details). From the GO enrichment analysis of each pairwise DEGs set built up using *ARAGNE* (Supplementary Material S1 and Supplementary Fig. S3), we show that genes upregulated in *pLAC17*-associated translatome were enriched for ribosomal and translation-related GO terms compared to *pVND6*-associated translatome (Supplementary Fig. S3A), and for vascular histogenesis terms compared to *pSWEET17*-associated translatome (Supplementary Fig. S3B). Up-regulated genes in *pSWEET17*-associated translatome were enriched with secondary cell wall biogenesis, xylan metabolism, and stress responses, including hypoxia, cold, and defense terms when compared to both *pLAC17*- and *pVND6*-associated translatomes (Supplementary Fig. S3C-D). In contrast, upregulated genes in *pVND6*-associated translatome were strongly enriched for photosynthesis and light-harvesting GO terms relative to both other populations, with fold enrichments exceeding 20 (Supplementary Fig. S3E-F). Together, these results suggest distinct functional identities of each translatome: *pLAC17*-associated cells are enriched for translational activity, *pSWEET17-*associated cells for cell wall synthesis and stress responses, and *pVND6*-associated cells for photosynthetic functions.

Further intersection analysis revealed 804 genes uniquely differentially expressed between the *pLAC17*- and *pVND6*-associated translatomes, 389 genes between the *pLAC17*-and *pSWEET17*-associated translatomes, and 247 genes between the *pSWEET17*- and *pVND6*-associated translatomes (Supplementary Fig. S2A, Supplementary Data Set 3). Furthermore, a set of 19 genes was found to be differentially expressed in all three pairwise comparisons simultaneously (Supplementary Fig. S2A-B and Supplementary Data Set 3). This intersection reflects genes with the strongest differential translational regulation across the three translatomes, with a clear expression gradient across the three translatomes (e.g., *pLAC17* > *pSWEET17* > *pVND6*, *pSWEET17* > *pLAC17* > *pVND6*) which confirms that each translatome contains subsets of genes with unique signatures across the three tissues that were analyzed (Supplementary Fig. S2B and Supplementary Data Set 3).

### Identification of translatome-enriched gene sets

Pairwise comparisons treat each contrast independently, meaning that a gene upregulated in *pLAC17-* vs. *pSWEET17-*associated translatomes is not necessarily upregulated in *pLAC17-* vs. *pVND6-*associated translatomes, preventing the direct identification of population-specific signatures. Therefore, we applied a directional intersection approach across the three pairwise comparisons to identify genes enriched in each translatome (see ‘Material and Methods’ section for more details). This technique allows the classification of genes according to their relative enrichment in each translatome (Supplementary Fig. S4; Supplementary Data Set 4). For example, a gene upregulated in the *pLAC17*-associated translatome compared to both the *pSWEET17*- and *pVND6*-associated translatomes was classified as enriched in the *pLAC17*-associated translatome. Using this approach, we identified 449 genes with clear translatome-specific enrichment, including 116 enriched in the *pLAC17*-associated translatome, 195 in the *pSWEET17*-associated translatome, and 138 in the *pVND6*-associated translatome (Supplementary Fig. S4 and Supplementary Data Set 4). The remaining 1,709 genes were initially excluded, as they were significant in at least one pairwise comparison but could not be unambiguously assigned to a single translatome. Nonetheless, to retain this information, we subsequently classified these genes based on their maximum expression level across translatomes (Supplementary Fig. S4 and Supplementary Data Set 4). This extended classification resulted in 843, 675, and 640 genes enriched in the *pLAC17*-, *pSWEET17*-, and *pVND6*-associated translatomes, respectively (Supplementary Fig. S4 and Supplementary Data Set 4). A new functional enrichment analysis was then performed using the lists of enriched genes identified in each translatome.

### Over-representation analysis of enriched genes in pVND6-associated translatome

Functional enrichment analysis of the enriched genes in *pVND6*-associated translatome identified 64 significantly enriched GO terms (Fig. 2 and Supplementary Table S2). Categories linked to cell wall organization and biogenesis (GO:0009832 and GO:0042546) and water or lignin transport processes (GO:0006833, GO:0015250) were overrepresented. In the context of cell wall biogenesis, it is notable that the *pVND6*-associated translatome includes genes involved in both primary and secondary cell wall synthesis. Specifically, genes such as *CELLULOSE SYNTHASE* (*CESA*) *2*, *CESA5*, *CESA6*, *XYLOGLUCAN ENDOTRANSGLUCOSYLASE/HYDROLAGE* (*XTH*) *31* and *XTH32*—primarily associated with primary cell wall formation and cell elongation—are present alongside genes like *FASCILIN-LIKE ARABINOOGALACTAN* (*FLA) 2*, *FLA9*, and *FLA11*, which are more directly linked to secondary cell wall synthesis (Persson et al. 2005, 2007; Desprez et al. 2007; Kaewthai et al. 2013; MacMillan et al. 2013). This suggests that different processes involved during primary and secondary cell wall deposition are represented. Interestingly the GO enrichment analysis of the *pVND6*-associated translatome also revealed a strong overrepresentation of categories related to carbohydrate metabolism and energy production. Terms associated with carbohydrate metabolic process (GO:0005975), glucose metabolic process (GO:0006006), polysaccharide metabolic process (GO:0005996), and cellular carbohydrate metabolic process (GO:0044262) were significantly enriched, together with processes linked to organic and primary metabolic pathways (GO:0016051, GO:0006082) (Fig. 2 and Supplementary Table S2). Consistent with the high energetic demand of secondary cell wall formation, we also observed enrichment in generation of precursor metabolites and energy (GO:0006091), oxidative phosphorylation and electron transport chain components (GO:0022900), proton transmembrane transport (GO:0015994), and photosynthesis-related categories including photosynthesis (GO:0015979), light harvesting (GO:0009765), and photosystem I and II assembly/function (GO:0009767, GO:0009768, GO:0009769) (Fig. 2 and Supplementary Table S2). In addition, multiple stress- and stimulus-responsive terms were enriched, such as response to heat (GO:0009416), response to abiotic stimulus (GO:0009266, GO:0009642, GO:0009644, GO:0009645), and response to hormone signaling pathways including auxin (GO:0009733) and other endogenous stimuli (GO:0009606) (Fig. 2 and Supplementary Table S2). Together, these enrichments indicate that the *pVND6*-associated translatome is enriched in genes involved of carbohydrate metabolism, bioenergetic pathways, photosynthetic functions, and stress/hormone-responsive processes, consistent with the extensive metabolic reprogramming required for xylem differentiation (Fig. 2).

**Figure 2.**
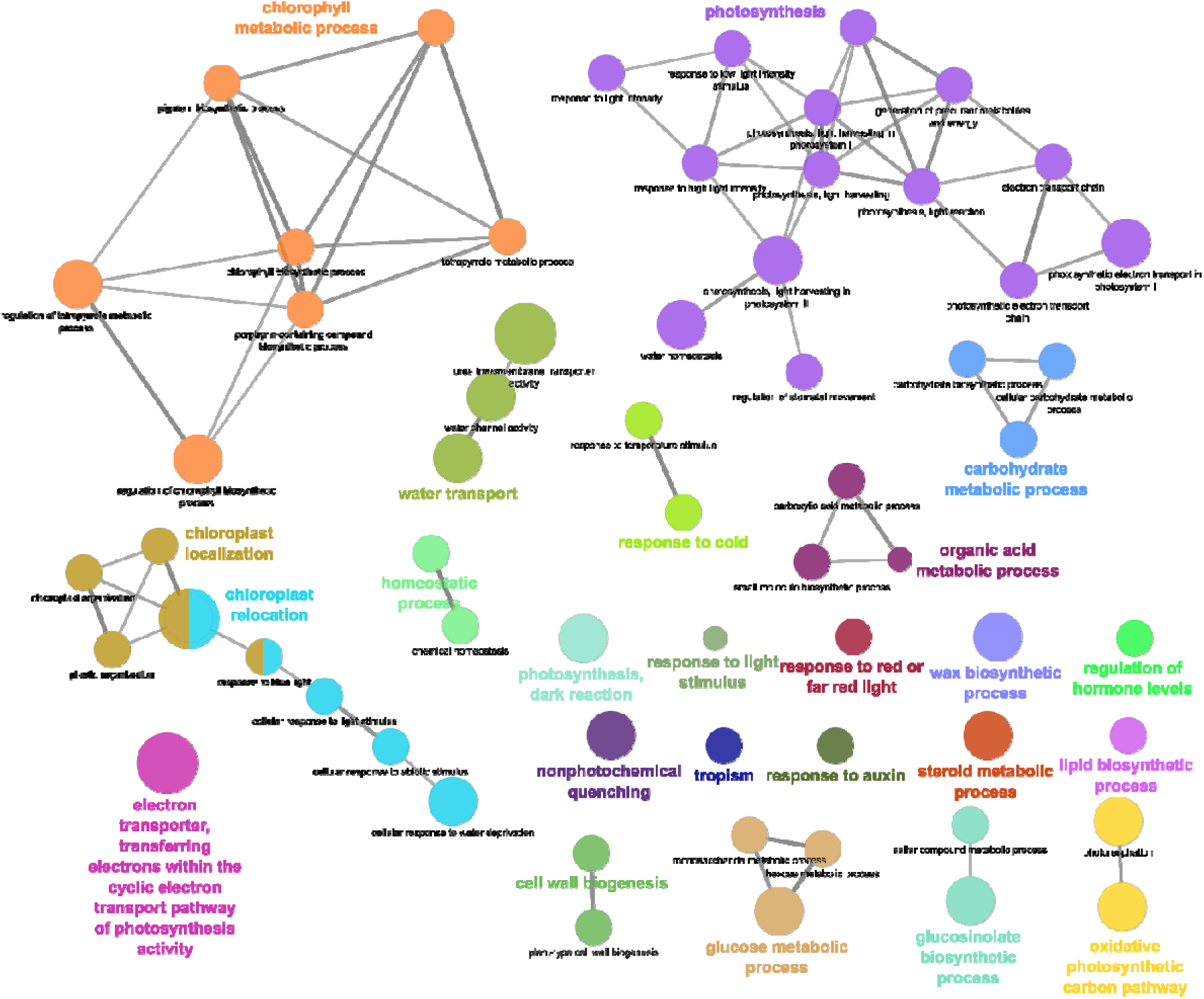
GO enrichment network of 640 enriched genes in *pVND6*-associated translatome. Gene Ontology (GO) enrichment analysis was performed on the set of genes identified as enriched in *pVND6*-associated translatome in comparison with *pLAC17*- and *pSWEET17*-associated gene sets. The list of genes is provided in Supplemental Data Set 4. Enrichment analysis was conducted using ClueGO and visualized in Cytoscape, as detailed in the *Materials and Methods*. Node size reflects the number of genes associated with each GO term in the TAIR11 background annotation and edges thickness is related to the KAPPA score. Only GO terms related to Biological Processes (BP) and for which a p-value lower than 0.05 are shown. The full list of enriched GO terms is available in Supplementary Table S2.

Especially, the strong enrichment in photosynthesis-related process points to an undifferentiated or very early stage of vessel development. To further assess this, we examined the expression of key cambial domain markers, including *PHLOEM INTERCALATED WITH XYLEM* (*PXY*) and *WUSCHEL RELATED HOMEOBOX 4* (*WOX4*) (proximal cambium/xylem) and *SMAX1-LIKE 5* (*SMXL5*) (distal cambium/phloem), which define distinct cambial domains in the Arabidopsis stem (Shi et al. 2021). While no significant change in *PXY* expression was observed across the translatome datasets (Supplementary Fig. S5A), *WOX4*—which typically mirrors *PXY* expression at the transcriptional level—was enriched in the *pLAC17*- and *pVND6*-associated translatomes relative to the *pSWEET17*-associated translatome (Supplementary Fig. S5B). Similarly, *SMXL5*, a marker of the distal cambium, was also enriched in the *pVND6*- and *pLAC17*-associated translatomes compared to *pSWEET17*-associated translatome (Supplementary Fig. S5C). Altogether, these results suggest that the *pVND6*-associated translatome represents a transitional cellular state, combining some cambial features with the activation of differentiation-related metabolic programs, thereby reflecting the progression toward xylem vessel differentiation.

### Over-representation analysis of genes enriched in pSWEET17-associated translatome

Gene ontology (GO) enrichment analysis of the genes enriched in *pSWEET17*-associated translatome revealed multiple terms related to plant secondary cell wall biogenesis and xylem histogenesis (e.g., GO:0009832, GO:0009834, GO:0071669, GO:0042546, GO:0010087, GO:2000652) as well as GO terms related to response to biotic and abiotic stresses (e.g., GO:1901700, GO:1901701, GO:1901698, GO:0010033, GO:0009266, GO:0009642, GO:0009644, GO:0009615, GO:0002237, GO:0002239) (Supplementary Fig. S6 and Supplementary Table S3). This suggests that, although the analysis focused on genes enriched in the *pSWEET17*-associated translatome relative to the other datasets—thereby highlighting features specific to xylem parenchyma cells, based on the observed expression pattern (Fig. 1), the dataset contain distinct sub-populations of cells including for example thin-walled xylem parenchyma cells, developing xylem vessels or thick-walled xylem parenchyma cells, undergoing distinct stages of SCW remodeling. Notably, the genes involved in the cell wall-related GO terms seems to be related to a later developmental stage than that represented in the *pVND6*-associated translatome.

To assess whether our dataset contains genes potentially expressed in thin-walled xylem parenchyma cells, or developing vessels we compared it with a recent single-cell RNA sequencing dataset from *Arabidopsis* roots undergoing secondary growth, which also includes both cell types (Lyu et al. 2025). Although our dataset reflects the translatome (*pSWEET17*-associated) and the other the transcriptome, and one focuses on stems while the other on roots, we anticipated identifying shared genes representing the xylem parenchyma signature across organs and regulatory levels. Among the 675 genes enriched in the *pSWEET17*-associated translatome (Supplementary Data Set 4), 607 were also detected in the single-cell RNA-seq dataset of roots (Supplementary Table S4). Of these, 53% (321 genes) exhibited maximal expression in xylem parenchyma cells (including mature, maturing, and young parenchyma), whereas the remaining 47% (286 genes) showed maximal expression during various stages of xylem vessel differentiation (Supplementary Table S4).

We then used the subset of *pSWEET17*-associated genes with peak expression in xylem parenchyma cells for a GO enrichment network analysis using CLUEGO app in Cytoscape (Fig. 3A and Supplementary Table S5). This network revealed that these genes, also expressed in root thin-walled xylem parenchyma cells, are strongly associated with biotic stress perception and downstream defense signaling, with notable contributions from hormone-mediated immunity pathways. Significantly enriched terms included responses to other organisms and bacteria (GO:0051707, GO:0009617), highlighting genes involved in the plant responses to abiotic and biotic stimuli. Multiple GO categories related to immune regulation—such as regulation of immune system process (GO:0002682), defense response, pathogen-associated molecular pattern signaling, and innate immune signaling—were overrepresented, reflecting different phases of the plant immune response (Fig. 3A and Supplementary Table S5). In addition, GO terms related to the plant responses to abiotic stress such as water deprivation (GO:0009415), hypoxia (GO:0001666), salt stress (GO: 0009651), osmotic stress (GO:0006970) and oxidative stress (GO:0006979) were also enriched. Consistently we observed enrichment for hormone-dependent signaling associated with immune responses, including jasmonic acid, salicylic acid, and ethylene responses, (GO:0009867, GO:0009723, GO:0009753, GO:0009863), suggesting a central role for hormone crosstalk in xylem parenchyma cells (Fig. 3A and Supplementary Table S5). Collectively, these GO terms indicate an enrichment in perception, signaling, and transcriptional regulation modules, in line with adaptation to biotic and abiotic stresses, in thin-walled xylem parenchyma cells.

**Figure 3.**
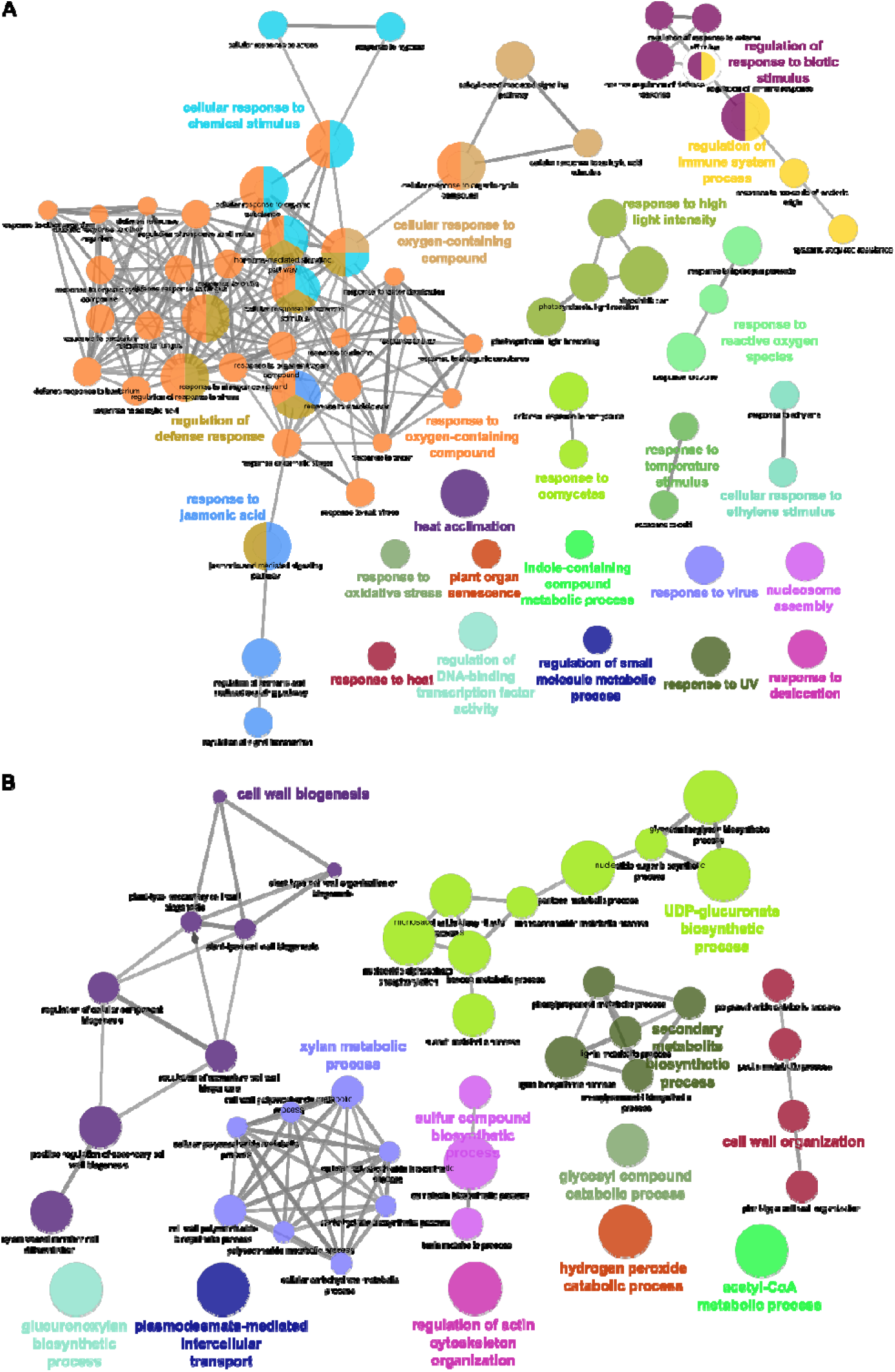
GO enrichment of the 607 enriched genes *pSWEET17*-associated stem translatome also present in the root xylem single-cell transcriptome. (A) GO enrichment network of 321 enriched genes in *pSWEET17*-associated stem translatome in common to the root xylem parenchyma single-cell transcriptome. (B) GO enrichment network of 286 enriched genes in *pSWEET17*-associated stem translatome in common to the root xylem vessels single-cell transcriptome. Enrichment was conducted using ClueGO and visualized in Cytoscape, as detailed in the *Materials and Methods*. Node size reflects the number of genes associated with each GO term in the TAIR11 background annotation and edges thickness is related to the KAPPA score. Only GO terms related to Biological Processes (BP) and for which a p-value lower than 0.05 are shown. The full list of genes used to perform the GO enrichment analysis can be found in Supplementary Table S4 and the list of enriched GO terms is available in Supplementary Table S5 and S6.

A similar analysis was performed on genes shared between the *pSWEET17*-associated translatome and those preferentially expressed in root xylem vessels during secondary growth (Fig. 3B and Supplementary Table S6). As expected, these genes were predominantly associated with cell wall construction and remodeling, tightly linked to phenolic secondary metabolism and primary carbohydrate pathways (Fig. 3B and Supplementary Table S6). Enrichment of cell wall biogenesis and plant-type cell wall organization terms (GO:0042546, GO:0071669) indicates active polysaccharide synthesis, deposition, and restructuring, consistent with growth- or stress-induced reinforcement of the extracellular matrix. GO terms related to carbohydrate and polysaccharide metabolism (GO:0005976, GO:0005996, GO:0005982, GO:0006024) further support engagement of primary carbon metabolism in providing precursors for structural polymers. Concurrent enrichment of phenylpropanoid and related aromatic compound biosynthetic processes (GO:0009698, GO:0009699, GO:0009808, GO:0009809, GO:0019321) suggests extensive lignin and wall-bound phenolic production, reflecting the reinforcement and chemical specialization of the cell wall represented in this subset of the *pSWEET17*-associated translatome. Considering that thick-walled xylem parenchyma cells have been described as lowly or even unlignified cells (Hoffmann et al. 2020), this subset likely reflect a later stage of xylem vessels differentiation.

### Over-representation analysis of genes enriched in pLAC17-associated translatome

The functional enrichment analysis of the genes specifically enriched in *pLAC17*-associated translatome revealed 64 significantly enriched GO terms (Fig. 4 and Supplementary Table S7). Among them, developmental terms such as secondary cell wall biogenesis (GO:0009834), plant-type cell wall organization (GO:0009832), vascular tissue development (GO:0003002), and xylem development (GO:0010089, GO:0009888) were significantly enriched as expected based on the expression domain of *pLAC17* mainly in interfascicular fibers (Fig. 4 and Supplementary Table S7). The enriched GO terms include key genes coding for fibers markers such as NST (NST1/At2g46770, SND1/NST3/NAC012/At1g32770 and NST2/NAC066/At3g61910) and MYB transcription factors associated with fiber development and secondary cell wall formation (MYB43/At5g16600, MYB52/At1g17950 and MYB69/At4g33450) (Ko et al. 2007; Schuetz et al. 2012; Taylor-Teeples et al. 2015; Geng et al. 2020; Smit et al. 2020). Interestingly, GO enrichment analysis of the *pLAC17*-associated translatome also revealed a marked overrepresentation of categories related to gene expression and macromolecule biosynthesis (Fig. 4 and Supplementary Table S7). Broad metabolic terms such as cellular nitrogen compound metabolic process (GO:0034641), organonitrogen compound metabolic process (GO:1901566), cellular macromolecule metabolic process (GO:0044260), and primary metabolic process (GO:0009059) were significantly enriched, indicating a global activation of anabolic pathways (Fig. 4 and Supplementary Table S7). Consistently, gene expression (GO:0010467), RNA processing (GO:0006364), mRNA splicing via spliceosome (GO:0000398), and translation (GO:0006412) were among the most significantly enriched categories. A strong signature of ribosome biogenesis and assembly was also detected, including ribosome biogenesis (GO:0042254), ribonucleoprotein complex biogenesis (GO:0022613), rRNA processing (GO:0000470, GO:0000463), and structural constituent of ribosome (GO:0002181). Terms linked to protein targeting and localization, such as protein targeting to mitochondrion (GO:0006605) and protein-containing complex assembly (GO:0043933), were enriched as well, together with chaperone-mediated protein folding (GO:0044085) and ubiquitin-dependent protein catabolic process (GO:0006518), highlighting an active protein quality control system (Fig. 4 and Supplementary Table S7). In addition, categories associated with cellular respiration and energy production were overrepresented, including generation of precursor metabolites and energy (GO:0006091), oxidative phosphorylation (GO:0006119), electron transport chain (GO:0022900), mitochondrial ATP synthesis coupled electron transport (GO:0042775), and tricarboxylic acid cycle-related processes (GO:0006120, GO:0006122). In summary, these results indicate that the *pLAC17*-associated translatome is characterized by translational capacity, ribosome production, mitochondrial bioenergetics, and cell wall-related developmental programs, consistent with the high metabolic and biosynthetic demands associated with lignification and secondary cell wall formation occurring primarily in the interfascicular fibers.

**Figure 4.**
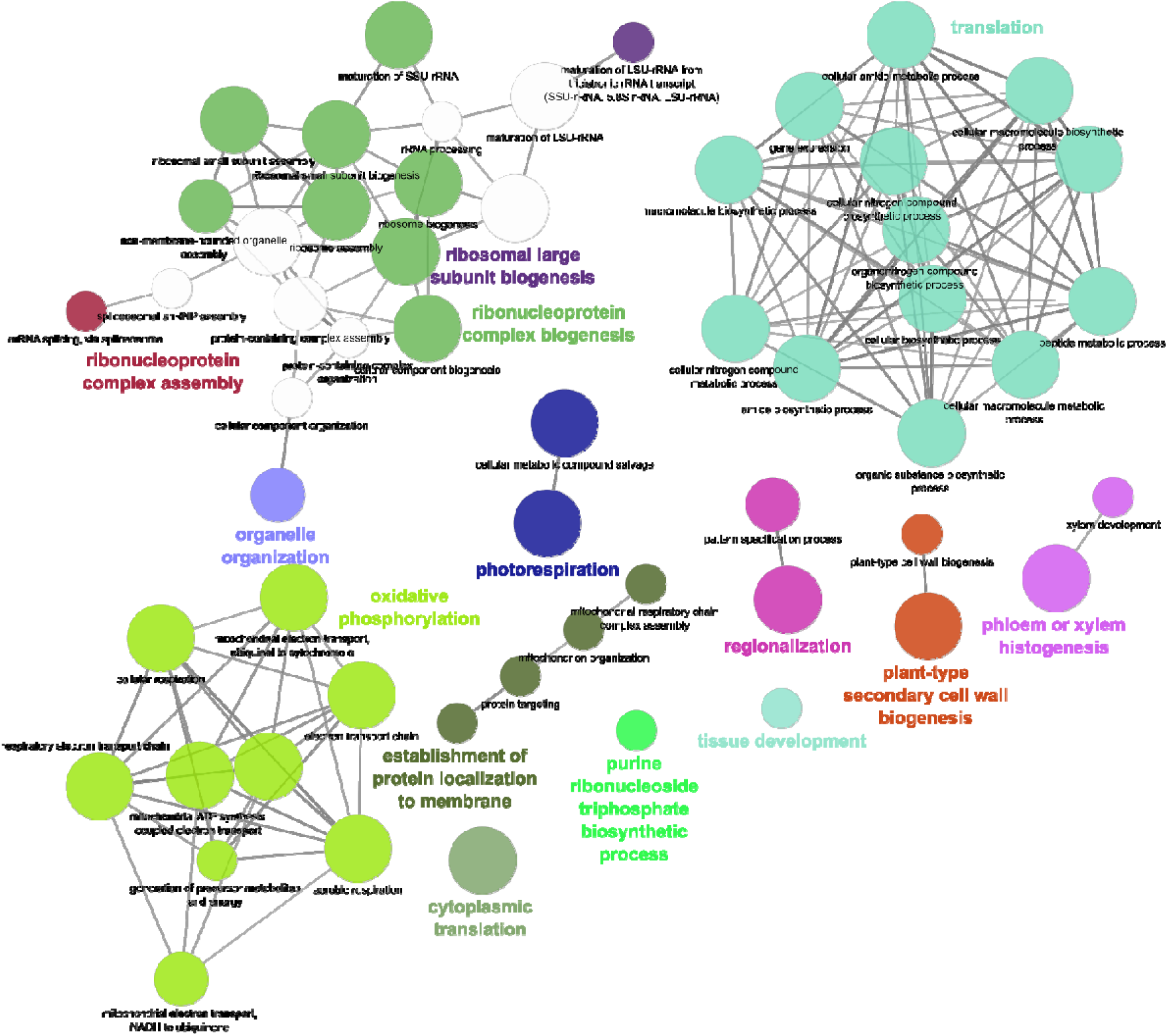
GO enrichment network of 843 enriched genes in *pLAC17*-associated translatome. Gene Ontology (GO) enrichment analysis was performed on the set of genes identified as enriched in *pLAC17*-associated translatome in comparison with *pVND6*- and *pSWEET17*-associated gene sets. The list of genes is provided in Supplemental Data Set 4. Enrichment analysis was conducted using ClueGO and visualized in Cytoscape, as detailed in the *Materials and Methods*. Node size reflects the number of genes associated with each GO term in the TAIR11 background annotation and edges thickness is related to the KAPPA score. Only GO terms related to Biological Processes (BP) and for which a p-value lower than 0.05 are shown. The full list of enriched GO terms is available in Supplementary Table S7.

### Sugar dynamics-related genes are actively translated across translatomes

Since carbohydrates metabolism was identified across the three translatome datasets, we asked whether our data could uncover new aspects of sugar dynamics across different cell types. To address this, we queried our translatome datasets using a curated list of 155 genes involved in sugar transport, metabolism, and signaling (Supplementary Table S8). Over 58% of these genes were actively translated across all three translatomes, although not necessarily differentially translated, including several with established roles in vascular development such as *FRUCTOKINASE 4* (*FRK4*), *FRUCTOKINASE 7* (*FRK7*) (Stein et al. 2017), coding for enzymes producing fructose-6-phosphate, *SWEET11* (Le Hir et al. 2015), *SWEET17* (Aubry et al. 2022), coding for sugar facilitators, and *SUCROSE SYNTHASE 1* (*SUS1*) and *SUS6* genes coding for enzymes involved in sucrose synthesis (Stein and Granot 2019).

To further refine this analysis, we compared our data with the transcriptomic atlas of inflorescence stem tissues generated by Shi et al. (2021), encompassing the transcriptome of xylem vessels and fibers. We found 84 common genes between our dataset and that of Shi et al. (2021), among which 59 genes were significantly differentially expressed in Shi et al. (2021) (Supplementary Table S9). Notably, more than half of these genes (34 out of 59) showed preferential expression in tissues associated with proximal cambium/xylem (*PXYpro:H4-GFP*), fibers (*NST3pro:H4-GFP*), and early vessel elements (*VND7pro:H4-GFP*), suggesting that they represent xylem-enriched isoforms (Supplementary Table S9). The remaining differentially expressed genes were mainly localized in starch sheath cells (*SCRpro:H4-GFP*). These included genes involved in sucrose metabolism (*VACUOLAR INVERTASE 1/VINV1*, *CELL WALL INVERTASE 3*/*CwINV3*), starch metabolism (*APD-GLUCOSE PYROPHOSPHORYLASE 1/ADG1*, *APD-GLUCOSE PYROPHOSPHORYLASE LARGE SUBUNIT 3/APL3*, *BETA-AMYLASE 1*/*BAM1*, *BETA-AMYLASE 3/BAM3*), sugar transport (*EARLY RESPONSE TO DEHYDRATION SIX-LIKE 3.08 /SUGAR TRANSPORTER ERD6*, *ESL3.12/ERDL11*, *ESL2.03/ERDL16*, *ESL3.13/SFP1/ERDL17*), and sugar signaling (*TREHALOSE-6-PHOSPHATE SYNTHASE (TPS) 5*, *TPS7*, *TPS9*, *TPS10*, and *TPS11*) (Supplementary Table S9).

Next, we explored whether our list of genes of interests (GOIs) related to sugar transport, metabolism or signaling (Fig. 5 and Supplementary Table S8) showed differential expression patterns in any of our translatome datasets by querying them in the *ARAGNE* app. This revealed that only *ESL3.04/ERDL13* was found to be enriched in the *pLAC17*-associated translatome (Fig. 5 and Supplementary Material S1). Furthermore, *UDP-GLUCOSE DEHYDROGENASE (UGD) 2*, *UGD3*, *UGD4*, *FRK1/2* and *FRK4/FRK7* were enriched in the *pSWEET17*-associated translatome (Fig. 5 and Supplementary Material S1). Finally, five genes coding for members of the tonoplast ESL/ERDL sugar transporter family (i.e., *ESL3.08/ERD6, ESL1.01/ERDL4*, *ESL3.06/ERDL2/ESL2*, *ESL2.03*/*ERDL16* and *ESL3.01/ERDL5*) and the *SWEET2* gene were found enriched in *pVND6*-associated translatome (Fig. 5 and Supplementary Material S1). The presence of these numerous tonoplast sugar transporters points to an important role of vacuole-cytosol sugar exchanges during early xylem development.

**Figure 5.**
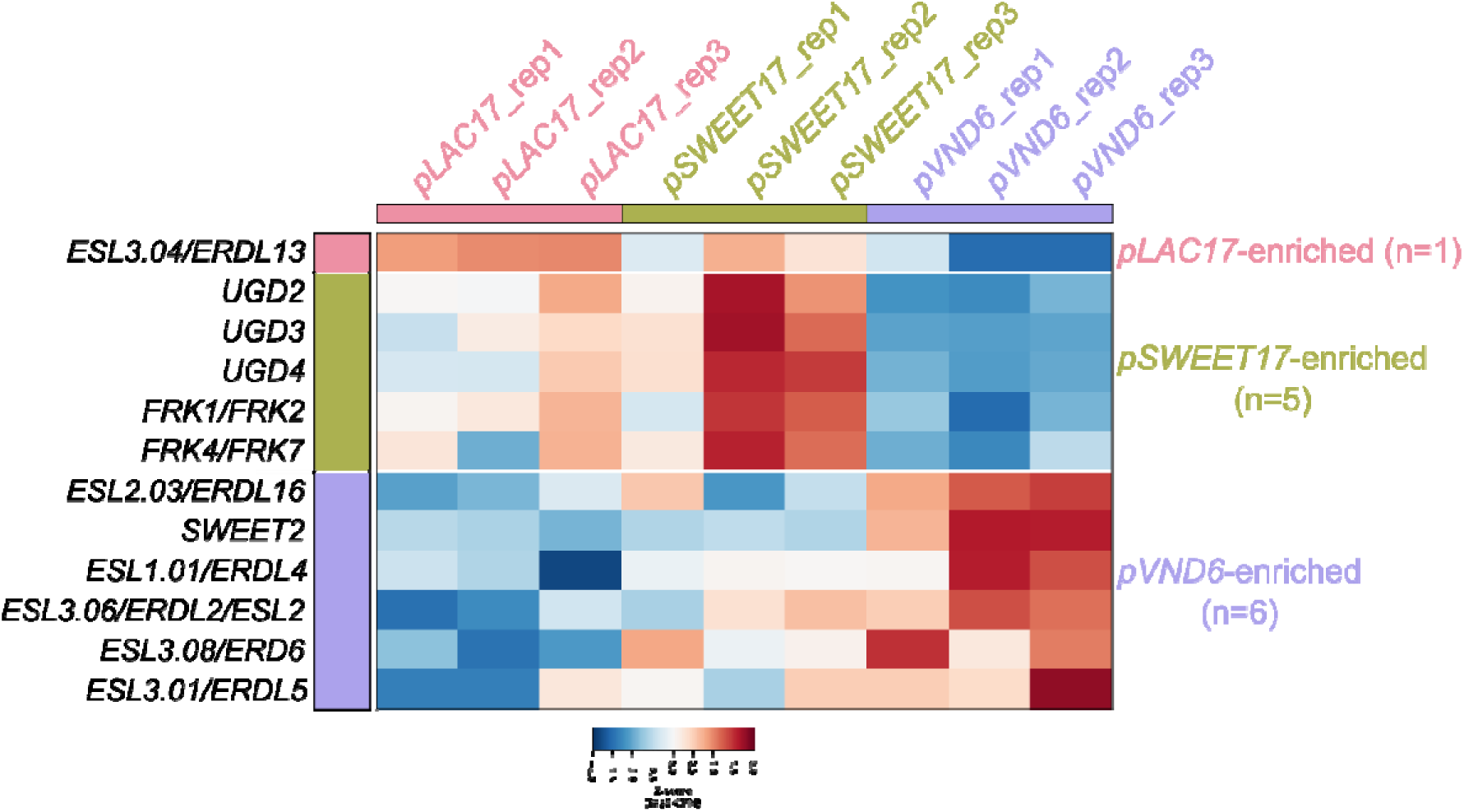
Heatmap of genes related to sugar transport, metabolism or signaling differentially enriched in each translatome. Color intensity represents the Z-score of log2-CPM values across the nine samples (three biological replicates per dataset). Genes are grouped by their assigned dataset and ordered by hierarchical clustering (Ward linkage, Euclidean distance) within each group. The list of genes queried can be found in Supplementary Table S8.

The presence of *SWEET2* is especially noteworthy, as we previously demonstrated that the two other tonoplast-localized SWEET transporters, *SWEET16* and *SWEET17*, play important roles in xylem development and secondary cell wall formation (Aubry et al. 2022). Therefore, using the *ARAGNE* app coupled with Cytoscape, we generated subnetworks around *SWEET2* for both pairwise comparison with *pVND6*-associated translatome (Supplementary Fig. S7 and Supplementary Table S10). Network analysis of the *SWEET2*-centered subnetwork revealed several highly connected and central genes potentially contributing to the structure and functional organization of the network. Among them, At1g12900/GAPA-2 (GLYCERALDEHYDE-3-PHOSPHATE DEHYDROGENASE A SUBUNIT 2) involved in glycolysis, displayed the highest connectivity (degree = 65) and betweenness centrality (3535.8), suggesting a major hub position within the network. Other highly central nodes included At1g49750, a LEUCINE-RICH REPEAT (LRR) family protein, At3g07870/F-BOX PROTEIN92/FBX92, and At1g15000/SERINE CARBOXYPEPTIDASE-LIKE 50 (SCPL50), all exhibiting high betweenness values (>1800) and moderate clustering coefficients (∼0.26–0.28), indicating potential roles as connectors between different functional modules. Several genes associated with metabolic and regulatory processes were also prominent in the network, including At1g20020/FERREDOXIN-NADP-REDUCTASE/FRN2, At3g25840/PRE-mRNA PROCESSING FACTOR 4 KINASE A/PRP4KA, At2g42160/BRAPS2 RING ZnF UBP DOMAIN-CONTAING PROTEIN 1/BRIZ1, and At5g55040/BROMODOMAIN-CONTAINING PROTEIN 13/BRD13. Within this subnetwork, *SWEET2* showed intermediate network properties, with a degree of 23, a betweenness centrality of 690.6, and a clustering coefficient of 0.41 (Supplementary Table S10). These values indicate that *SWEET2* is moderately connected and embedded within a locally cohesive neighborhood while maintaining connections with several other nodes in the network. Moreover, these results indicate that the *SWEET2*-associated subnetwork is composed of genes involved in diverse cellular functions, including metabolism (At3g08590/2,3-BIPHOSPHOGLYCERATE-INDEPENDENT PHOSPHOGLYCERATE MUTASE/iPGAM2), signaling (At4g20360/RAB GTPASE HOMOLOG 8A/RABE1C), and plastid-related processes (At1g08510/FATTY ACYL-ACP THIOESTERASES B/FATB, At1g42550/PLASTID IMPAIRED1/PMI1 and At3g15095/HIGH CHLOROPHYLL FLUORESCENCE 243/HCF243), and that *SWEET2* occupies an intermediate but structurally relevant position within this network. Since *SWEET2* was found to be enriched in *pVND6*-associated translatome, we further investigated its role in xylem development, and more broadly, the contribution of tonoplast SWEET sugar transporters to the development of the stem vascular system.

### SWEET2 is expressed in the stem vascular system and is required for stem radial growth

SWEET2 has previously been characterized as a glucose-specific transporter, predominantly expressed in roots and involved in plant defense against *Pythium* infection (Chen et al. 2015). In this study, we show that *SWEET2* is also expressed in the Arabidopsis inflorescence stem. Its expression is significantly enriched in the *pVND6*-associated translatome, although it is also present in the other two translatomes (Fig. 6A). To further confirm its expression pattern, we used a reporter line from Chen et al. (2015), which expresses a translational GUS fusion with the C-terminus of SWEET2 under the control of its native promoter in a wild-type background. GUS staining revealed that, at the bottom part of the stem, SWEET2 is expressed in all xylem tissues, including developing metaxylem vessel elements, xylem parenchyma cells, and both fascicular and interfascicular fibers which is consistent with our TRAP-seq data (Fig. 6B-C). Additional expression was also observed in the phloem and pith (Fig. 6B-C).

**Figure 6.**
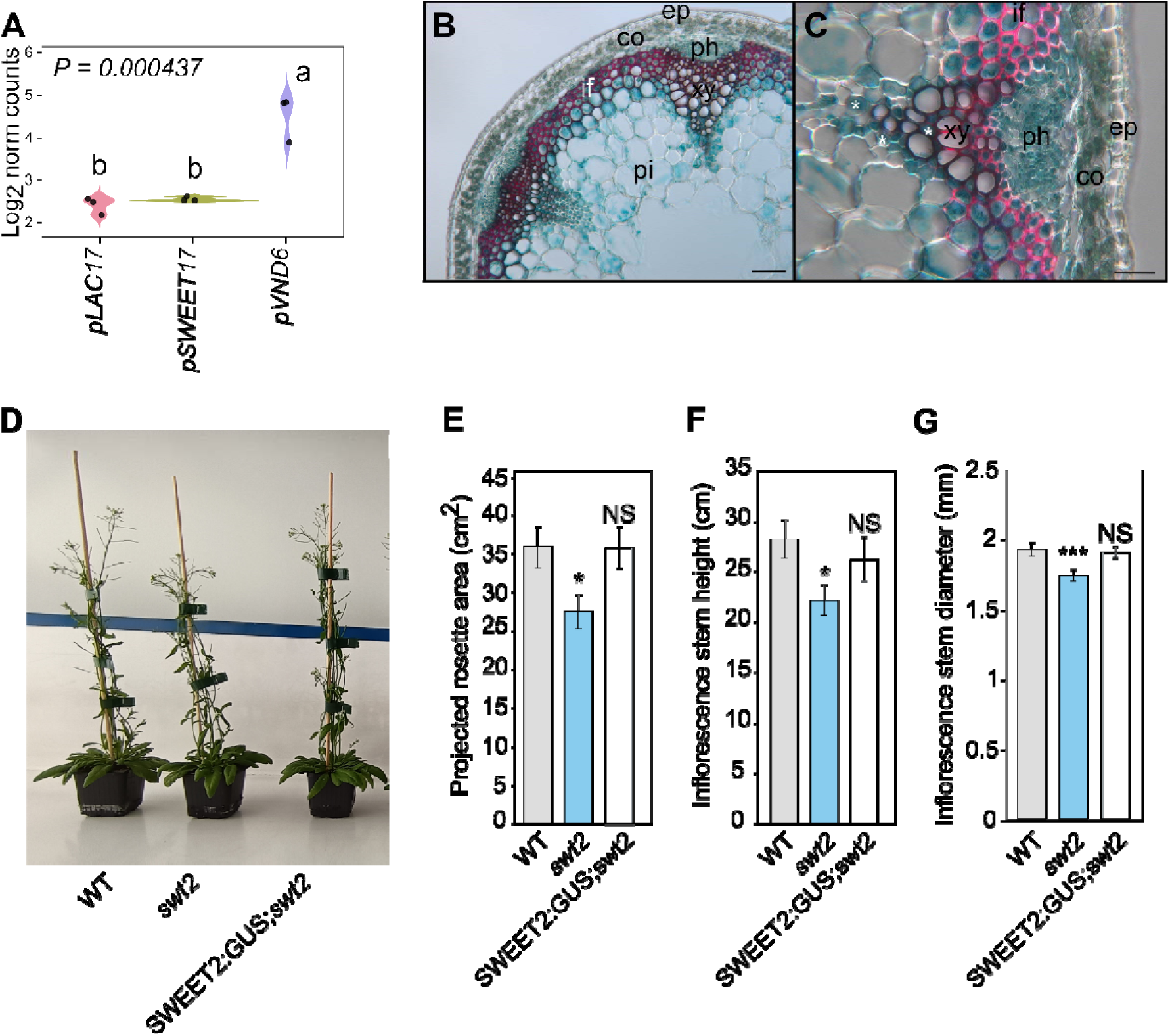
SWEET2 is expressed in the stem vascular system and its disruption induced defect in radial stem growth. (A) Violin plot showing log_2_-transformed normalized gene reads counts of the *SWEET2* among the different translatomes. Each black dot corresponds to a biological replicate (n = 3 biological replicates). A one-way ANOVA combined with Tukey’s comparison post-test has been made to compare the different translatomes. The different letters indicate significant differences. The *P*-value for the comparison between the different translatomes is indicated on the graph. (B-C) Histochemical analysis of GUS activity in lines expressing SWEET2-GUS fusion protein driven by the SWEET2 native promoter in sections taken in the inflorescence stem of 45-day-old plants. Asterisks points to xylem parenchyma cells. Lignin is colored pink after phloroglucinol staining. ep, epidermis; co, cortex; if, interfascicular fibers; ph, phloem; pi, pith; xy, xylem. Scale bar = 100 µm (B) or 50 µm (C). (D) Barplot showing the inflorescence stem of the single *swt2* and its corresponding complemented line SWEET2:GUS;*swt2*. Means ± SE (n ≥ 10 plants for each genotype coming from two independents experiments). A student t-test was performed to compare the different line to the wild-type plants. *** *P*<0.001, ** *P*<0.01, NS: not significant.

To investigate the functional role of SWEET2 in stem development, we analyzed the *sweet2-1* (*swt2*) mutant previously described by Chen et al. (2015). Under our growth conditions, *swt2* plants exhibited significantly reduced overall growth compared to wild-type, including smaller rosettes and decreased inflorescence stem height and diameter (Fig. 6D-G). To confirm that these growth defects were due to the loss of *SWEET2* function, we generated complemented lines expressing pSWEET2:SWEET2-GUS in its mutant background. Rosette size, stem height and stem diameter were fully restored to wild-type levels, confirming the direct role of SWEET2 in the overall plant growth including radial stem growth (Fig. 6E-G).

### Non-additive effects of *SWEET2*, *SWEET16*, and *SWEET17* mutations on xylem development

To date, three tonoplast-localized SWEET sugar transporters have been characterized in *Arabidopsis*: SWEET2, SWEET16, and SWEET17 (For review see Keller et al. 2026). SWEET2 has been proposed to function as a vacuolar glucose transporter (Chen et al. 2015), whereas SWEET17 acts as a fructose facilitator (Chardon et al. 2013). SWEET16 exhibits broader substrate specificity and is capable of transporting sucrose, glucose, and fructose (Klemens et al. 2013). In addition, SWEET16 and SWEET17 are expressed in the stem vascular system and have been implicated in xylem development and secondary cell wall formation (Aubry et al., 2022). To further elucidate the role of SWEET2 in stem development, and to assess the combined impact of mutations affecting all three tonoplast SWEET transporters, we generated and functionally characterized a series of single and higher-order mutant lines. This included the single mutants *swt2*, *swt16*, and *swt17*; the double mutants *swt2swt16* (*swt2;16*), *swt2swt17* (*swt2;17*), and *swt16swt17* (*swt16;17*); as well as the triple mutant *swt2swt16swt17* (*swt2;16;17*). As part of two additional independent cultures, the projected rosette area, the inflorescence stem height and diameter were measured across all mutant lines (Fig. 7A and Supplementary Fig. S8). We confirmed that *swt2* mutant displays smaller rosette, inflorescence stem height and diameter (Fig. 7A and Supplementary Fig. S8). Moreover, the projected rosette area of the *swt2;17* double mutant is also significantly smaller than that of wild-type plants and resembled to the *swt2* single mutant line (Supplementary Fig. S8A). On the other hand, the rosette size of the other mutant lines did not significantly differ from wild-type plants (Supplementary Fig. S8A). Regarding the stem height, none of the mutant lines (except the *swt2* mutant) display a significant change compared to wild-type plants (Supplementary Fig. S8B). Interestingly all mutant lines developed significantly thinner stems than wild-type plants (Fig. 7A). This phenotype has already been previously reported for the *swt16;17* double mutant (Aubry et al. 2022).

**Figure 7.**
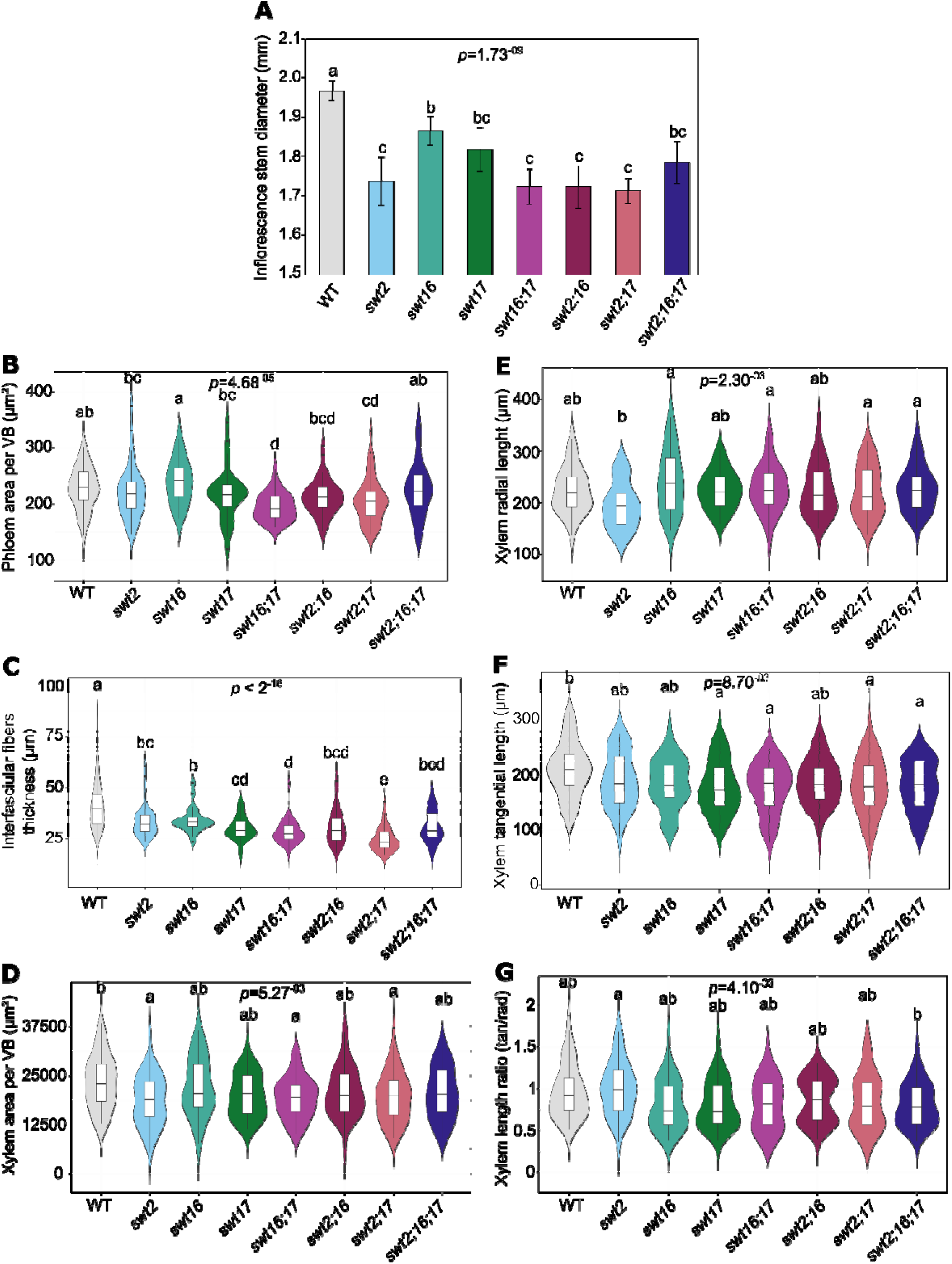
Loss of tonoplast SWEET transporters affects fascicular and interfascicular tissue development. (A) Inflorescence stem diameter in wild-type (WT), single (*swt2*, *swt16*, *swt17*), double (*swt16;17*, *swt2;16*, *swt2;17*), and triple (*swt2;16;17*) mutant lines. Bars show least-square means ± SE from two independent experiments (n ≥ 10 plants per genotype). (B–G) Violin plots quantifying anatomical features related to the vascular system: (B) phloem area per vascular bundle (VB), (C) interfascicular fiber thickness, (D) xylem area per VB, (E) xylem radial length, (F) xylem tangential length, and (G) xylem length ratio (as tangential/radial axes ratio). Values are from individual VBs (WT, n = 47; *swt2*, n = 46; *swt16*, n = 39; *swt17*, n = 41; *swt16;17*, n = 44; *swt2;16*, n = 51; *swt2;17*, n = 62; *swt2;16;17*, n = 53) across two independent experiments (n ≥ 10 plants per genotype). Statistical significance was assessed using one-way ANOVA followed by Tukey’s post hoc test. Different letters indicate significant differences (P < 0.05). Exact *P*-values are shown on the plots.

Notably, the stem of the *swt2;16* double mutant resembled that of the *swt2* single mutant, highlighting a predominant role of *SWEET2* (Fig. 7A). Since SWEET2 specifically mediates glucose transport, these results suggest that vacuole-to-cytosol glucose transport by SWEET2 is critical for stem radial growth. Likewise, the *swt2;17* double mutant closely mirrored the *swt2* phenotype (Fig. 7A), further supporting a central role of *SWEET2* in this process. Interestingly, the *swt2;16;17* triple mutant did not exhibit a stronger phenotype than the single mutants, indicating a non-additive effect in stem diameter reduction (Fig. 7A).

To better understand the anatomical basis of these changes, we performed a morphometric analysis of the vascular system by quantifying the average areas of phloem and xylem tissues within vascular bundles and measuring the thickness of the interfascicular fibers (Fig. 7B–D). Additionally, xylem radial and tangential lengths were measured as indicators of cell proliferation and expansion, respectively, and their ratio was calculated as a proxy for xylem shape (Fig. 7E–G) (Wang et al. 2019). Phloem area did not differ significantly between wild type and single mutants (Fig. 7B). In contrast, it was significantly reduced in the *swt16;17* and *swt2;17* double mutants, whereas the *swt2;16* line showed only a downward trend without statistical significance (Fig. 7B). Strikingly, in the *swt2;16;17* triple mutant, the phloem area was comparable to wild type, suggesting partial compensation (Fig. 7B). The interfascicular fiber thickness was significantly reduced in all mutant lines (Fig. 7C). Among the single mutants, *swt17* showed the strongest reduction, and the *swt16;17* double mutant closely resembled this phenotype. The *swt2;16* double mutant was not further affected compared to its single mutants, whereas the *swt2;17* double mutant displayed the most pronounced reduction, indicating an additive effect. Interestingly, the triple mutant differed significantly from the *swt2;17* line but not from the single mutants, suggesting that loss of *SWEET16* partially counteracts the *swt2;17* phenotype (Fig. 7C). In terms of xylem development, the *swt2*, *swt16;17* (as also reported in Aubry et al., 2022), and *swt2;17* mutants exhibited significantly smaller xylem areas compared to wild type, while the other mutants were unaffected (Fig. 7D). Radial length (a proxy for cell number) did not differ across genotypes (Fig. 7E). By contrast, tangential length (a proxy for cell expansion) was significantly reduced in *swt17*, *swt16;17*, *swt2;17*, and *swt2;16;17* mutants, indicating smaller xylem cells in these lines (Fig. 7F). No significant differences in xylem shape (radial/tangential length ratio) were observed relative to wild type, except between *swt2* and *swt2;16;17*, suggesting that adding *SWEET16* and *SWEET17* mutations modifies xylem shape even if the resulting phenotype is not significantly different from wild-type plants (Fig. 7G).

In summary, these findings indicate that the reduced stem diameter in *sweet* mutants arises from defects in both fascicular and interfascicular tissues. Moreover, they point out a new role for *SWEET2* in interfascicular fibers and xylem development. They also raise an important role of glucose and fructose vacuole-to-cytosol transport mediated by both SWEET2 and SWEET17 during these processes.

### Organ-specific effects of *SWEET* transporter mutations on sugar metabolism

Next, we analyzed the sugar status in stems and rosette leaves of the different mutant lines (Fig. 8). In stems, the *swt2* mutant accumulated significantly higher levels of both glucose and fructose compared to wild type (Fig. 8A–B), whereas sucrose content was unchanged and starch content showed a non-significant upward trend (Fig. 8C–D). Interestingly, although *swt16* and *swt17* mutants also tended to accumulate more glucose, the addition of mutations led to a significant reduction of stem glucose, as observed in the *swt2;17* and *swt2;16;17* lines (Fig. 8A). Similarly, stem sucrose content was significantly reduced only in the double and triple mutants (Fig. 8C). Starch levels were consistently higher in all mutant lines, with significant increases detected only in *swt2;16* and *swt2;17*, while the remaining mutants displayed intermediate phenotypes (Fig. 8D). In rosette leaves, the *swt2* mutant also accumulated markedly higher glucose than wild type (Fig. 8E). Strikingly, in the *swt2;16;17* triple mutant, glucose levels were restored to wild-type values, suggesting compensatory metabolic adjustments when vacuolar glucose export is strongly impaired (Fig. 8E). Fructose content was significantly increased in *swt2;17* and *swt2;16;17*, while *swt2* and *swt16* did not differ from wild type, and *swt17* showed a non-significant upward trend (Fig. 8F). No significant changes in leaf sucrose or starch content were observed across genotypes (Fig. 8G–H).

**Figure 8.**
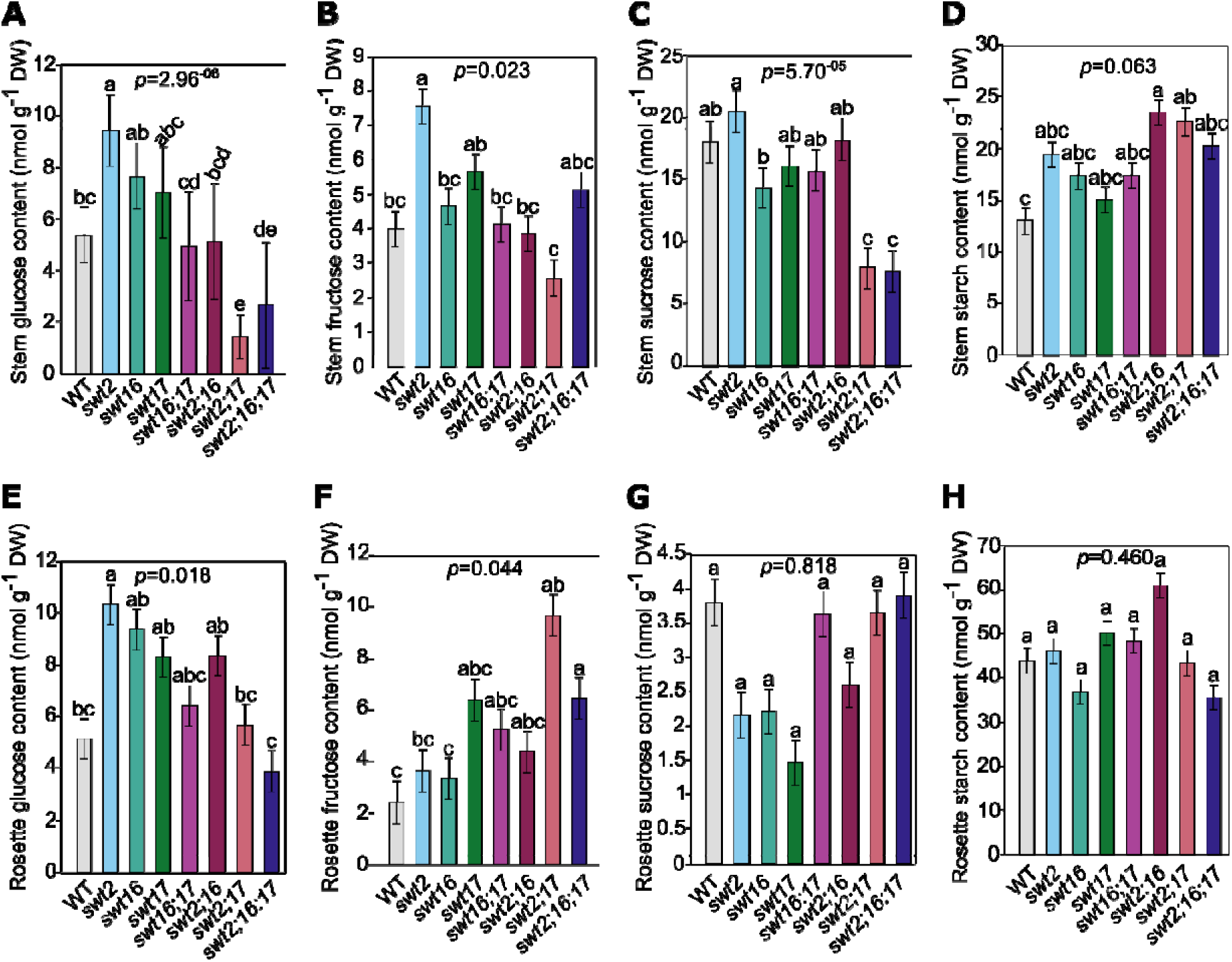
Mutations in tonoplastic *SWEET* transporters lead to differential changes in sugar content in both stem and rosettes leaves. (A-H) Barplots showing the glucose (A and E), fructose (B and F), sucrose (C and G) and starch (D and H) contents in inflorescence stem (A-D) and the rosette leaves (E-H). Means ± SD are shown (4 ≥ n ≤ 6 for each genotype). A one-way ANOVA combined with Duncan’s comparison post-test has been made to compare the different genotypes. The different letters indicate significant differences. The *P*-values for the comparison between the different genotypes are indicated on each graph.

Together, these results indicate that mutations in genes coding for SWEET tonoplast transporters trigger organ-specific metabolic responses. In particular, the reduced stem diameter phenotype of the *swt2* mutant could be associated with altered hexose partitioning.

### Xylem and interfascicular cell wall composition is impaired in the different mutant lines

In this work, we show that SWEET2 is expressed in both xylem tissue and interfascicular fibers (Fig. 6). Moreover, our previous studies identified that *SWEET16* and *SWEET17* are required for normal secondary cell wall composition (Aubry et al. 2022). Therefore, we explore whether loss of *SWEET2*, alone or in combination with *SWEET16* and/or *SWEET17*, affects the secondary cell wall composition, and whether such effects are tissue dependent. To this end, we performed Fourier-transformed infrared spectroscopy (FTIR) in inflorescence stem cross-sections to analyze the xylem and interfascicular fibers secondary cell wall composition (Fig. 9 and Supplementary Fig. S9).

**Figure 9.**
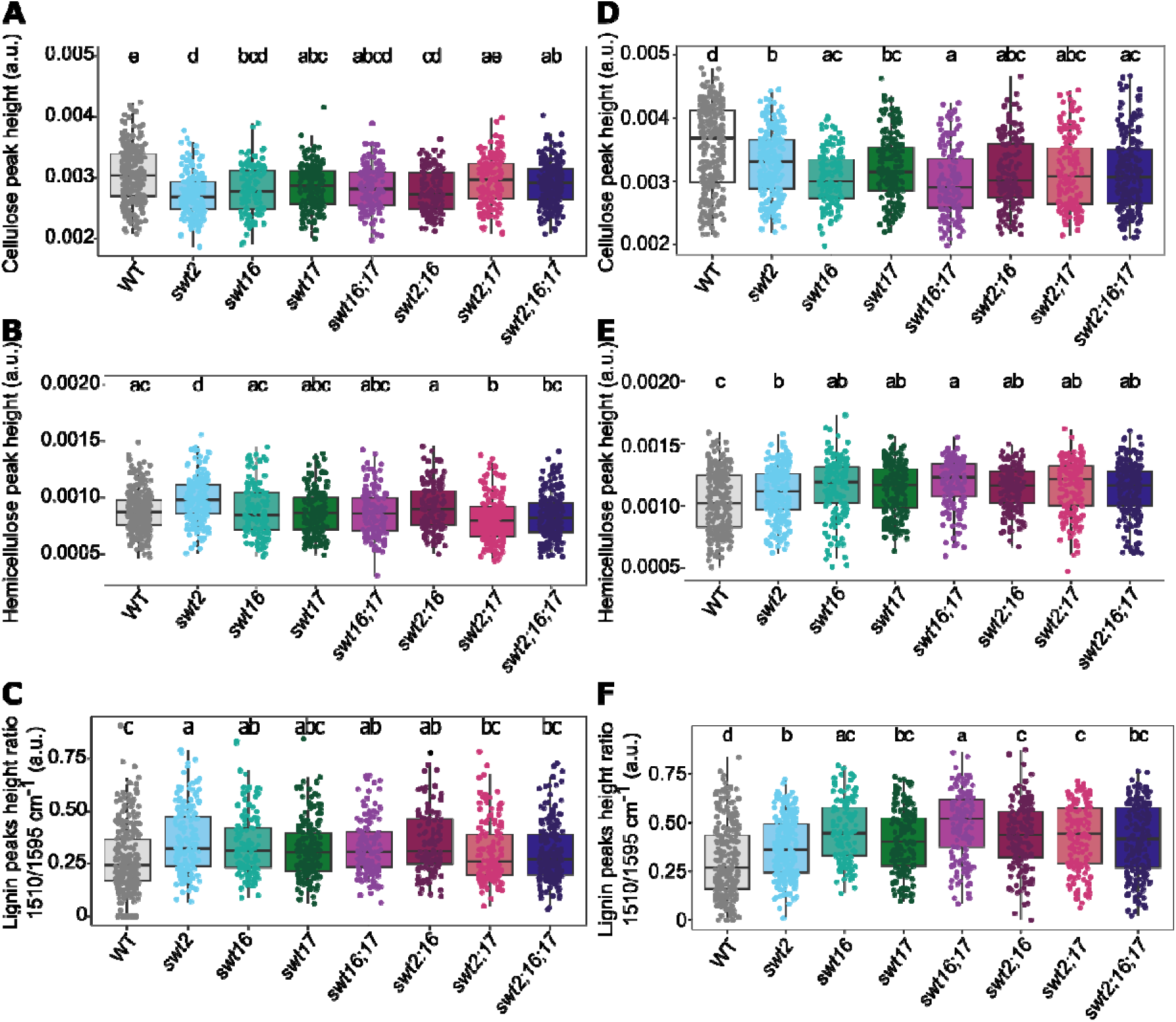
Impact mutations of tonoplast *SWEET* on secondary cell wall composition of xylem and interfascicular fibers. FTIR spectra were acquired on xylem tissue (A-C) and on interfascicular fiber (D-F) from sections of the basal part of the inflorescence stem. All spectra were baseline-corrected and area-normalized in the range 1800-800 cm^-1^. From these spectra, the cellulose (C–O vibration band at 1,050 cm^-1^) (A and D), hemicellulose (C–O and C–C bond stretching at 1,740 cm^-1^) (B and E) peak height and lignin peak height ratio (1,510/1,595 cm^-1^) (C and F) were measured and plotted as boxplots. The lines represent median values, the tops and bottoms of the boxes represent the first and third quartiles, respectively, and the ends of the whiskers represent maximum and minimum data points. The boxplots represent values (shown as colored dots) from 272/253 (xylem/interfascicular fibers), 186/249, 127/143, 182/188, 139/146, 133/168, 168/167, and 205/208 spectra from the wild-type, *swt2*, *swt16*, *swt17*, *swt16;17*, *swt2;16*, *swt2;17* and *swt2;16;17* plants, respectively, obtained from 9-12 plants for each genotype grown in two independent cultures. A one-way analysis of variance combined with the Tukey’s comparison post hoc test was performed. Values marked with the same letter were not significantly different from each other, whereas different letters indicate significant differences (P C 0.05).

In xylem tissues, the average FTIR spectra of all mutant lines revealed multiple differences relative to the wild type within the cellulose, hemicellulose, and lignin fingerprint regions (Supplementary Fig. S9A). Consistently, *t*-value spectra showed several significant positive and negative peaks at wavenumbers associated with cellulosic and hemicellulosic polysaccharides, including 898 cmC¹, the 995–1120 cmC¹ region, 1187 cmC¹, and the 1373–1485 cmC¹ region (KacCuráková et al. 2000; Åkerholm and Salmén 2001; Lahlali et al. 2015) (Supplementary Fig. S9C). Notably, a decrease at 1429 cmC¹ together with an increase at 898 cmC¹ is typically associated with reduced cellulose crystallinity (Nelson and O’Connor 1964). This opposite pattern was observed in the *swt2;17* and *swt2;16;17* mutant lines, suggesting increased cellulose crystallinity in these genotypes (Supplementary Fig. S9C). In contrast, the *swt2* single mutant displayed increased absorbance at both 898 and 1429–1430 cmC¹, indicating a distinct alteration in cellulose organization. Altogether, these data point to defects in cellulose crystallinity or organization across multiple *sweet* mutant lines. In addition, several genotypes (*swt16*, *swt17*, *swt16;17*, and *swt2;16*) showed a significant reduction in the cellulose-associated peak at 1050 cmC¹, consistent with previous observations in the *swt16;17* mutant (Aubry et al. 2022) (Fig. 9A).

Regarding hemicelluloses, all mutant lines exhibited a significant increase in the 1369 cmC¹ band, which is associated with C–H deformation of CHC acetyl groups in hemicelluloses and acetylated cellulose (Mohebby 2008) (Supplementary Fig. S9C). Moreover, the 1740 cmC¹ band, characteristic of acetylated xylan (Gou et al. 2008), was significantly increased in the *swt2* single mutant, indicating enhanced xylan acetylation (Supplementary Fig. S9C). In contrast, this band was significantly reduced in the *swt2;17* and *swt2;16;17* mutants, suggesting decreased xylan acetylation in these backgrounds. Quantification of the 1740 cmC¹ peak height confirmed these opposing trends between *swt2* and the *swt2;17* and *swt2;16;17* lines (Fig. 9B). Interestingly, the *swt2;16* double mutant closely resembled the *swt16* single mutant and did not differ significantly from the wild type, indicating that loss of *SWEET2* alone is not sufficient to explain the xylan acetylation phenotypes observed (Fig. 9B). Nevertheless, these results support an important role for SWEET2 in matrix polysaccharide deposition in xylem secondary walls.

Finally, the lignin peak height ratio (1510/1595 cmC¹) was significantly increased in the *swt2*, *swt16*, *swt16;17*, and *swt2;16* mutant lines (Fig. 9C), suggesting a relative enrichment in guaiacyl (G) units compared with syringyl (S) units, which may contribute to altered SCW rigidity.

In interfascicular fibers, the overall spectral profiles, *t*-value spectra, and cellulose-associated peak height changes were largely similar to those observed in xylem tissues (Supplementary Fig. S9B and D). However, more genotypes exhibited significant differences in the 1740 cmC¹ band and in the 1510/1595 cmC¹ lignin ratio (Fig. 9E-F). In contrast to xylem tissues, the *swt16*, *swt17*, *swt16;17*, *swt2;16*, and *swt2;16;17* mutants showed a significant increase in acetylated xylan relative to the wild type (Fig. 9E). Notably, the *swt2;17* mutant displayed an opposite behavior compared with xylem, with a significant increase in the 1740 cmC¹ peak height in interfascicular fibers (Fig. 9E). Similarly, the 1510/1595 cmC¹ lignin ratio was significantly altered in the *swt17*, *swt2;17*, and *swt2;16;17* mutants in interfascicular fibers, whereas no such differences were detected in xylem tissues (Fig. 9F). Thus, these results indicate that mutations in tonoplast-localized SWEET sugar transporters, among which SWEET2 seems predominant, exert stronger effects on hemicellulose and lignin composition in interfascicular fibers than in xylem tissues.

## DISCUSSION

As the main contributor to lifetime carbon gain in *Arabidopsis* (Earley et al. 2009), the inflorescence stem is a key photosynthetic and transport organ required for developing fruits. It also exhibits strong developmental gradients along both radial and longitudinal axes, encompassing vascular and interfascicular cell types at distinct developmental stages. The inflorescence stem has therefore been used as a model organ to investigate the molecular networks underlying vascular development and secondary cell wall formation, notably through bulk and tissue-specific transcriptomic approaches (Vanholme et al. 2012; Hall and Ellis 2013; Shi et al. 2021). However, transcriptomic analyses, while capable of identifying lowly expressed regulators such as transcription factors and transporters—particularly with the increased resolution provided by single-cell approaches—do not reflect their translational status. Indeed, transcript abundance does not always correlate with protein synthesis, as some transcripts may be subject to translational regulation that enhances or, conversely, strongly limits their translation. Moreover, transcript-level analyses can overlook molecular actors whose regulation primarily occurs at the translational level, such as sugar-responsive genes (Gamm et al. 2014). To gain deeper insight into the molecular networks operating in xylem vessels, parenchyma cells, and interfascicular fibers, we employed a TRAP-seq approach. This strategy enables tissue-type-specific profiling of the translatome, providing new insights into the functional roles of these tissue types and revealing previously unrecognized molecular actors that warrant further investigation.

### Cell type-specific translational regulation of a core vascular gene set

The 19 genes differentially expressed across all three pairwise translatome comparisons represent the most robustly cell type-regulated fraction of the vascular translatome, with each gene displaying a quantitative translational hierarchy that distinguishes interfascicular fibers, xylem parenchyma cells, and very early xylem vessel differentiation from one another simultaneously (Supplementary Fig. S2B). Interestingly, their functional annotation is coherent with the known biology of each cell type.

In *pLAC17*-associated translatome illustrating expression in interfascicular fibers, the highest translational activity of two ribosomal protein-encoding genes (*RIBOSOMAL PROTEIN L41*/*RPL41*, *RIBOSOMAL PROTEIN ES30Z*/*RPS30A*) is consistent with the intense biosynthetic demand required for secondary cell wall deposition. This process has recently been shown to involve post-transcriptional regulation by Musashi-type RNA-binding proteins (Kairouani et al. 2023). The enrichment of a DOF transcription factor further suggests translational prioritization of regulators linked to carbon metabolism and cell wall biosynthesis (Le Hir and Bellini 2013; Zou and Sun 2023).

In *pSWEET17*-associated translatome (likely reflecting xylem parenchyma cells), the dominant translational signature is related to metabolic versatility and stress response. For instance, the *GALACTINOL SYNTHASE 2* (*GOLS2*), which catalyzes the first committed step in raffinose family oligosaccharides (RFOs) biosynthesis (Sengupta et al. 2015), is consistent with the role of xylem parenchyma cells in carbohydrate storage as proposed in trees (Secchi et al. 2016). Beyond acting as compatible solutes, RFOs also function as hydroxyl radical scavengers (Nishizawa et al. 2008), suggesting that xylem parenchyma cells may help buffer the oxidative microenvironment generated during lignification of adjacent vessel elements. The co-enrichment of the *ARABIDOPSIS THALIANA DROUGHT-INDUCED 8/ATDI8,* a dehydrin (Gosti et al. 1995; Méndez-Gómez et al. 2024), the *CALMODULIN LIKE 10* (*CLM10*/*CaBP-22*) a calmodulin-like EF-hand calcium-binding protein (Day et al. 2002), and the *GLYCINE-RICH PROTEINS 3S* (*GRP3S*) (Czolpinska and Rurek 2018) supports the idea that these cells maintain a constitutive stress-buffering program, consistent, likely contributing to xylem hydraulic stability (Secchi et al. 2016). Similarly, the presence of the *TIR-NS8*, a TIR domain-containing protein (At1g72910) (Nasim et al. 2020) and the *ARABIDOPSIS ORTHOLOG OF SUGAR BEET HS1 PRO-1 2*/*HSPRO2* suggests cell type-specific immune competence, potentially reflecting exposure to pathogen-derived signals within the xylem sap. The enrichment of *GLUCURONOXYLAN 4-O METHYLTRANFERASE 2*/*GXM2* (Yuan et al. 2014) further indicates a role in modifying in neighboring cell walls, consistent with the known intercellular coordination of cell wall biosynthesis in vascular tissues (Smith et al. 2013, 2017a).

Finally, very early xylem vessel differentiation (*pVND6*-associated translatome) is marked by the enrichment of the *AUXILIN-LIKE 1* (*AUL1*), an AUX/IAA transcriptional repressor. This aligns with the central role of auxin signaling in xylem specification (Smetana et al. 2019; Xu et al. 2019), and suggest that fine-tuning of auxin responses is regulated at the translational level during early differentiation.

Together, these genes define a coherent, cell type-resolved translational program that mirrors the distinct physiological identities of fibers, xylem parenchyma, and nascent vessel elements, and highlights translational regulation as a key layer of cell type specification in the plant vascular system.

### Early metaxylem differentiation integrates cell wall remodeling with primary metabolism

VND6, a NAC transcription factor, is a key regulator of metaxylem differentiation (Kubo et al. 2005; Zhong et al. 2008; Shi et al. 2021). Accordingly, the *pVND6*-associated translatome is enriched in genes involved in both primary and secondary cell wall biosynthesis (Fig. 2; Supplementary Table S2). Primary wall CESAs (CESA2, CESA5, CESA6), which function during cell expansion (Desprez et al. 2007; Persson et al. 2007), are enriched alongside secondary wall-associated genes, reflecting the known sequential action of distinct cellulose synthase complexes during xylem differentiation (Watanabe et al. 2018). Additional enrichment of *XTH* genes involved in wall remodeling and *FLA* genes associated with secondary wall formation (Kaewthai et al. 2013; MacMillan et al. 2013; Ma et al. 2022) indicates that the cells correspond to an early transitional stage, during which primary-to-secondary wall remodeling is actively occurring.

Some cells may also represent states between cambial initials and xylem precursors, as suggested by the presence of the cambial markers *WOX4* and *SMXL5* (Supplementary Fig. S5). This may account for the enrichment of photosynthesis- and metabolism-related genes (Fig. 2), which is consistent with previous studies highlighting the importance of primary metabolism during xylem differentiation (Ohtani et al. 2016; Verbančič et al. 2018). In particular, Ohtani et al. (2016) demonstrated that protoxylem differentiation involves a major reprogramming of primary metabolism, redirecting carbon flux toward the synthesis of cell wall polysaccharides and lignin monomers. Our results also further support that early xylem cells retain active plastids, supporting energy production and biosynthetic fluxes required for secondary wall formation, as observed in wood species across different biomes (Schmitz et al. 2012). Time-resolved transcriptomic and metabolomic analyses of xylogenesis have also shown that primary metabolism and organelle coordination—particularly involving chloroplasts and mitochondria—are prominent during early developmental stages and decline as secondary wall deposition and cell death programs intensify (Ohtani et al. 2016; Pinard et al. 2019). Consistently, lignin-deficient mutants show increased expression of chloroplast-associated proteins (Vanholme et al. 2012), pointing to a functional link between plastid activity and cell wall biosynthesis. Furthermore, tonoplast sugar transporters, such as ESL/ERDL family members and SWEET2, are enriched in the *pVND6*-associated translatome, suggesting a coordination of carbohydrate fluxes between the cytosol, vacuole, and plastids.

A subset of the *pSWEET17*-associated translatome likely represents later differentiation stages, retaining enrichment in carbohydrate metabolism and transport functions such as members of the fructokinases family implicated in vascular development (Stein et al. 2017); the enrichment of BAM1 (At3g23920), a chloroplastic enzyme involved in starch metabolism, and USUALLY MULTIPLE ACIDS MOVE IN AND OUT TRANSPORTERS 42 (UMAMIT42), a putative amino acid transporter localized to the tonoplast and Golgi/endosome (Zhao et al. 2021). Altogether, these data support a model in which early metaxylem differentiation integrates cell wall remodeling with active primary metabolism and inter-organelle coordination.

### Stem xylem parenchyma cells as hubs for metabolic integration and stress responses

Xylem parenchyma cells are key contributors to storage, metabolite exchange, and stress resilience, although their molecular characterization remains limited especially in herbaceous species (Barcelo 2005; Plavcová et al. 2013; Smith et al. 2013; Secchi et al. 2016; Słupianek et al. 2021). Recent single-cell analyses have begun to resolve their transcriptional dynamics in roots undergoing secondary growth (Lyu et al. 2025). Integrating these data with the *pSWEET17*-associated translatome identifies a core gene set consistently expressed in thin-walled xylem parenchyma cells (Fig. 3A; Supplementary Table S5).

Gene ontology analysis reveals strong enrichment in responses to biotic and abiotic stimuli, including pathogens, water deficit, osmotic stress, and temperature, together with strong enrichment in hormone signaling pathways (notably abscisic acid, jasmonic acid, salicylic acid, and ethylene) (Fig. 3A and Supplementary Table S5). This suggests that xylem parenchyma cells function as active sensory and signaling hubs rather than passive storage compartments. Moreover, the enrichment of immune-related processes, including systemic acquired resistance and responses to microbial elicitors (e.g., chitin and bacterial molecules), supports an unanticipated role for these cells in vascular immunity. This is further reinforced by the presence of key regulatory hubs such as WRKY (e.g., WRKY23, WRKY33, WRKY40, WRKY48, WRKY53), AP2/ERF (e.g., ETHYLENE RESPONSE FACTOR (ERF)4, ERF61, ERF109, ERF105, ERF104, ERF2, ERF11, ERF59, RELATED TO AP2 (RAP2) 4/RAP2.4, RAP2.1, and C-REPEAT-BINDING FACTOR 4/CBF4/DREB1D), and MYC2 transcription factors, as well as components of reactive oxygen species (ROS) and redox signaling pathways, indicating tight coupling between stress perception and downstream responses. The co-occurence of SNF1-RELATED PROTEIN KINASE 2 (SnRK2) kinases and AP2/ERF factors like RELATED TO ABI3/RAV1/EDF4 or ERF104 further support a role in integrating stress signaling pathways in response to biotic and abiotic stresses (Bethke et al. 2009; Feng et al. 2014). Concurrent enrichment of water deprivation, hypoxia, and oxidative stress responses also points to a role in buffering environmental fluctuations within the vascular tissue. This dataset thus offers a foundation for further elucidating how SnRK-mediated phosphorylation may modulate AP2/ERF activity in the context of environmental constraints, potentially revealing new mechanisms by which plants integrate environmental cues into xylem parenchyma-specific transcriptional responses. Together, these data support a model in which xylem parenchyma cells may integrate environmental, hormonal, and metabolic signals to coordinate vascular adaptation, defense, and homeostasis.

### Ribosome biogenesis and alternative splicing are linked to interfascicular fibers development

The *pLAC17*-associated translatome shows strong enrichment in ribosomal proteins, indicating intense protein synthesis during interfascicular fiber development (Fig. 4). At first glance, this observation may appear counterintuitive, since *LAC17* expression peaks at late stages of fiber development, just prior to programmed cell death (Berthet et al. 2011). However, previous studies in poplar have shown that ribosomal proteins are tightly regulated during wood formation, being either upregulated in tension wood relative to normal wood (Bygdell et al. 2017) or downregulated across xylem development (Obudulu et al. 2016). One possible explanation is that during secondary cell wall formation, cells must produce large amounts of proteins required for monolignol biosynthesis, lignin polymerization, and polysaccharide deposition, which would demand increased ribosome biogenesis and translational capacity. Moreover, even though fiber cells are ultimately destined to die, they may undergo a phase of hyperactivity—a “final burst” of biosynthesis—before shutdown. Interestingly, in mammalian systems ribosomes themselves have been implicated in apoptosis induction (Sinha et al. 2024), raising the possibility that ribosome dynamics could also contribute to programmed cell death in plants. The enrichment of spliceosome-related genes (Fig. 4) further suggests an important role for alternative splicing during interfascicular fiber development. Interestingly, evidence has emerged for a role of the spliceosome in xylem development in both herbaceous and woody plants (Ohtani et al. 2008; Chen et al. 2020). Our results support the idea that alternative splicing also contributes to the later stages of interfascicular fiber development. They also raise the intriguing possibility that spliceosome-mediated regulation could participate in inducing programmed cell death, as it has been described in plant–pathogen interactions (Wu et al. 2023).

### SWEET2: a central regulator in xylem and interfascicular fibers development

Our analysis identifies SWEET2, a tonoplast-localized glucose transporter, as a key regulator of stem development despites its absence or low detectability in conventional transcriptomic datasets (For review, see Dinant and Le Hir 2022; Lee et al. 2025). Nonetheless, tissue-specific RNA-seq revealed steady-state *SWEET2* expression across stem vascular cell types (Shi et al. 2021). This discrepancy likely reflects the preferential expression of *SWEET2* in vascular-associated subpopulations that represent only a minor fraction of total stem cells, causing transcript levels to fall below detection thresholds in single-cell atlases – or unexpected transcriptional or translational regulations depending on the cell type. Tissue-enriched RNA-seq, by integrating transcript abundance across vascular tissues, is therefore better suited to detect such restricted expression patterns. Consistent with this interpretation, our translatome datasets show that *SWEET2* is primarily enriched in early xylem precursor cells (*pVND6*-associated translatome), while translational GUS fusion assays revealed broader protein accumulation throughout the inflorescence stem (Fig. 5). Functional analyses further demonstrated that SWEET2 is required for normal stem development, with *swt2* mutants exhibiting reduced stem radial growth, a phenotype fully rescued in complemented lines (Fig. 6).

To place SWEET2 function in the context of SWEET-mediated vacuolar sugar transport, we further analyzed mutant combinations involving the three tonoplast-localized SWEET transporters expressed in stems: SWEET2, SWEET16, and SWEET17 (Aubry et al. 2022 and this work). These analyses revealed a clear functional hierarchy. SWEET2 emerged as the dominant regulator of stem radial growth, as all genotypes carrying the *swt2* allele—whether single, double, or triple mutants—displayed similarly reduced stem diameters (Fig. 7). The absence of additive phenotypes in higher-order mutants indicates that SWEET2 plays a rate-limiting, non-redundant role during stem thickening.

SWEET2 was previously characterized as a vacuolar glucose importer based on heterologous expression and plant experiments conducted under glucose-excess (Chen et al. 2015). In that study, *sweet2* mutants displayed reduced levels of several soluble sugars in leaves. By contrast, we observed a significant accumulation of glucose and fructose in both stems and rosette leaves of *swt2* mutants (Fig. 7). A key methodological difference between the two studies lies in growth conditions: Chen et al. grew plants under short-day conditions before shifting to long days to induce flowering, whereas our plants were grown continuously under long-day conditions from germination. Because photoperiod strongly influences carbon allocation and vacuolar buffering strategies in Arabidopsis (Sulpice et al. 2014), these contrasting sugar phenotypes likely reflect distinct carbon regimes. Indeed, under fluctuating or carbon-limiting conditions, SWEET2 likely functions primarily as a vacuolar glucose importer, sequestering excess cytosolic sugars during recovery from carbon starvation. Under continuous long-day conditions, however, the absence of *SWEET2* appears to impair vacuolar–cytosolic sugar homeostasis, leading to whole-organ hexose accumulation. We therefore propose that SWEET2 may also contribute to glucose export from the vacuole when cytosolic demand is high, consistent with the bidirectional, gradient-driven transport mode of SWEET proteins (Xue et al. 2022).

At the cellular level, *SWEET2* translation is enriched in early stage of xylem vessel differentiation relative to xylem parenchyma cells and interfascicular fibers (Fig. 6), suggesting cell-type-specific functions. At the stem level, the accumulation of soluble sugars in *swt2* mutant line points to reduced cytosolic sugar utilization, supporting a role for SWEET2 in the rapid adjustment of intracellular hexose levels (Fig. 10). In the context of its expression in early differentiating xylem vessel and interfascicular fibers—derived from interfascicular parenchyma cells (Mazur et al. 2014)—together with reduced stem growth and altered secondary wall composition in *swt2* mutants, suggests a role in supporting local UDP-glucose production and polysaccharide biosynthesis (Fig. 10). Overall, *SWEET2* disruption compromises intracellular sugar buffering and indirectly limits efficient carbon use for secondary cell wall formation.

**Figure 10.**
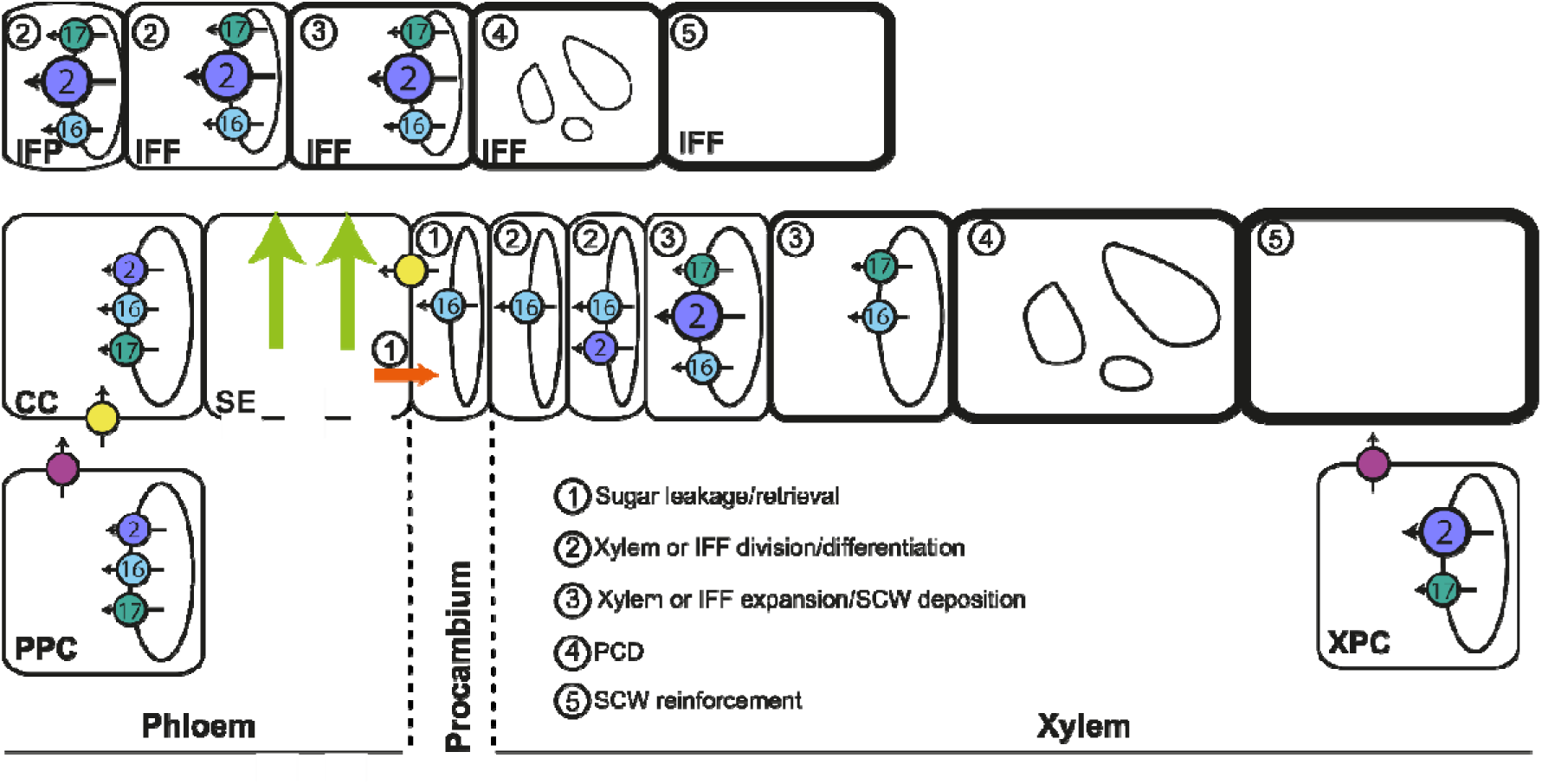
Model for the role of tonoplast SWEET transporters in vascular system development in Arabidopsis stems. This updated model integrates the results obtained in this study on SWEET2 with previously published data on SWEET16, SWEET17, SWEET11, SWEET12, and SUCROSE-PROTON SYMPORTER 2 (SUC2) (Truernit and Sauer 1995; Chen et al. 2012; Gould et al. 2012; Le Hir et al. 2015; Aubry et al. 2022). In phloem tissues, sugar exchanges between the cytosol and the vacuole in companion cells (CC) and/or phloem parenchyma cells (PPC) are regulated by the three tonoplast transporters SWEET2, SWEET16 and SWEET17. Cytosolic sucrose and hexoses in PPC are exported to the apoplast by SWEET11 and SWEET12 (fuchsia circles) and subsequently imported into CC by SUC2 (yellow circles), before entering sieve elements (SE) and being transported over long distances (light green arrows). A fraction of these sugars leaks from SE, likely through plasmodesmata (orange arrow), supplying axial sinks such as procambium and xylem, while another fraction is retrieved into SE, mainly via SUC2 (1). At the cambium–xylem boundary, soluble sugars are likely released from the vacuole into the cytosol by SWEET16 (light blue) to sustain xylem cell division (2). During secondary cell wall (SCW) deposition, which requires high cytosolic sugar availability (3), vacuolar sugar pools are probably mobilized through the combined action of SWEET2, SWEET16 and/or SWEET17. Following programmed cell death (PCD) and vacuole disintegration (4), SCW reinforcement continues (5), implying sustained sugar demand. At this stage, sugars stored in the vacuoles of xylem parenchyma cells (including xylary and axial xylem parenchyma) are likely released by SWEET2 and SWEET17 and exported to the apoplast by SWEET11 and SWEET12. In parallel, sugars stored in the vacuoles of interfascicular parenchyma cells (IFP) are exported by the three tonoplast SWEET transporters to support interfascicular fiber (IFF) development and SCW formation. To emphasize the central role of SWEET2 in xylem and interfascicular fibers development, the transporter is depicted with a bigger icon than SWEET16 and SWEET17.

At the system level, the topology of the *SWEET2*-centered subnetwork suggests that *SWEET2* is embedded within a regulatory context linking carbon metabolism, plastid-associated functions, and cellular signaling. Several highly central nodes in the network are associated with primary metabolism or plastid activity, including *GAPA-2*, a key enzyme of the Calvin–Benson cycle, and *FRN2*, which participates in photosynthetic electron transfer through the ferredoxin–NADP reductase system (Marri et al. 2005; Mulo 2011). The presence of these genes among the most connected and central nodes suggests that the *SWEET2* subnetwork may be functionally associated with pathways related to photosynthetic carbon assimilation and redox metabolism. Moreover, the presence of two genes coding for proteins belonging to the NAD(P)-binding Rossmann-fold superfamily protein, namely VEP1 and SDRD, involved in cytosolic steroid metabolism and the peroxisomal step of fatty acids metabolism, respectively (Jun et al. 2002; Herl et al. 2009; Quan et al. 2013) further suggest that tonoplast sugar transport mediated by SWEET2 is coordinated with lipid metabolism. Moreover, VEP1 has been shown to be involved in vascular strand development in Arabidopsis linking steroid metabolism and vascular development (Jun et al. 2002). In addition, the network includes genes coding for multiple regulatory proteins, such as the F-box protein FBX92 involved in cell division (Baute et al. 2017); a RING-type ubiquitin ligase BRIZ1 involved in early plant growth (Hsia and Callis 2010) and PRP4KA a spliceosome-associated kinase involved in alternative splicing (Kanno et al. 2018). Moreover, *SCPL50*, a member of the serine carboxypeptidase-like family, whose members function either in protein processing or as acyltransferases involved in specialized metabolism (Fraser et al. 2005) is also found the *SWEET2*-centered network. SCPL50 has recently been shown to regulate β-glucosidase 25 signaling during phosphate starvation by cleaving its ER-retention motif (Cho et al. 2025), but its presence in the Arabidopsis vacuolar proteome (Carter et al. 2004) suggests additional tonoplast-associated functions. Whether these proteins participate in SWEET2 regulation through alternative splicing, ubiquitination or proteolytic processing remains an open question, but their co-expression highlight the utility of network-based approaches for identifying candidate regulators of vacuolar sugar transport. Taken together, the network structure supports the hypothesis that SWEET2 participates in a broader regulatory module linking sugar transport with carbon metabolism, organelles activity, and signaling processes, further highlighting a role for vacuole-to-cytosol sugar exchanges in the integration of metabolic and regulatory cues within the cell.

In contrast to SWEET2, SWEET16 and SWEET17 exert more subtle, tissue-specific effects. While their single mutants showed little impact on overall stem growth, they displayed pronounced defects in interfascicular fiber thickness, xylem cell expansion, and secondary wall composition (Fig. 10). SWEET17, in particular, emerged as an important regulator of fiber differentiation, with reduced xylem tangential length, thinner interfascicular fibers, and altered lignin composition in *swt17* mutants. These observations reinforce the role of vacuolar fructose export in maintaining osmotic balance and supporting cell expansion and wall deposition (Aubry et al. 2022; Lu et al. 2025). The partial suppression or inversion of several traits in the triple mutant further indicates metabolic compensation and plasticity, arguing against simple functional redundancy among tonoplast SWEET transporters.

Taken together, our data support a model in which SWEET2 acts as a central regulator of stem radial growth by ensuring cytosolic hexoses availability for secondary wall biosynthesis, while SWEET16 and SWEET17 fine-tune tissue-specific aspects of cell expansion and wall composition (Fig. 10). These findings further underscore the importance of subcellular sugar partitioning and sugar identity—beyond total sugar availability—in shaping vascular development.

## MATERIALS AND METHODS

### Gene cloning, plasmid construction and transformation

Constructs used for TRAP-seq were generated by Gateway cloning. The sequences of the promoters of *LAC17*, *VND6* and *SWEET17* were first amplified using specific primers (Supplementary Table S11) (Kubo et al. 2005; Berthet et al. 2011; Aubry et al. 2022). Taking advantage of restriction sites generated by PCR, the purified PCR fragments were cloned into a pENTR modified donor vector. These donor vectors thus created were used to transfer, by LR recombination (LR clonase, ThermoFisher Scientific) the fragments into the destination vectors pGATA:HF-RPL18 (Mustroph et al. 2009). To generate GUS fusions constructs, the sequences of *LAC17pro-RPL18*, *VND6pro-RPL18* and *SWEET17pro-RPL18* were amplified using upstream specific primers (Supplementary Table S11) and RPL18 reverse primer without stop codon. The amplified fragments were then cloned into a pENTR modified vector, then introduced by LR recombination into pMDC162 (Curtis and Grossniklaus 2003) generating promoters-RPL18-GUS constructs. Destination vectors created this way were analyzed by sequencing to check the correct reading frame of the fusion. For plant transformation, binary vectors were introduced into *Agrobacterium tumefaciens* C58pMP90 (Koncz and Schell 1986) and plants were transformed by floral dip (Clough and Bent 1998). Transformants were selected on kanamycin (50 mg/ml) and homozygous T3 seeds were used for the analysis.

### Translating ribosome affinity purification and RNA extraction

Plants were grown in growth chamber conditions (Aubry et al. 2022) and sampled after 45 days after sowing (DAS). The siliques, flowers and lateral stems were removed and only the main stem was kept and frozen in liquid nitrogen. The stem was considered as a whole getting therefore information from different xylem developmental stages. At the sampling stage, the stem is still undergoing elongation but most of the internodes are not elongating anymore. Twenty-four stems per line were pooled and three independent cultures were performed. Frozen stem were ground using mortar and pestle and homogenized in ice-cold polysome extraction buffer and processed as described in Leal et al. (2022). After polysomes elution, the total RNA was extracted using the TRIzol reagent (Thermo Fisher Scientific, 15595-026, https://www.thermofisher.com) and treated with DNase I, RNase free (Thermo Fisher, EN0521, https://www.thermofisher.com) according to the manufacturer’s instructions. RNA concentration and purity were determined using Bioanalyzer 2100 (Agilent Technologies, Santa Clare, CA, USA). Samples RNA integrity number (RINs) were ranging from 6.9 to 8.7 and processed for library construction. The RNA samples were kept in −80°C until use.

### Library preparation, RNA sequencing, read mapping, data normalization and differential gene expression analysis

One ng of RNA produced previously were used as templates for first-strand cDNA synthesis of mRNA sequences close to the 3′end using QuantSeq 3′mRNA-Seq Library Prep Kit REV (Lexogen). The kit allows the generation of Illumina-compatible libraries from polyadenylated RNA and only generates one fragment per transcript (Moll et al. 2014). The libraries were quality checked (QC) using high sensitivity DNA assay with Bioanalyzer (Agilent). A total of 10 cDNA libraries were prepared for the three transgenic lines from three to four biological replicates. Libraries that passed QC were sequenced on the Illumina NextSeq 500 in single-end (SE) mode with 75 bases. Read quality was assessed using FastQC (https://www.bioinformatics.babraham.ac.uk/projects/fastqc/). Between 5.3 and 11.3M reads per sample were generated. UMIs were removed and appended to the read identifier with the extract command of UMI-tools (v1.0.1, Smith et al. 2017b). Reads where any UMI base quality score falls below 10 were removed. Low-quality tails, poly(A)read-through and adapter contamination were trimmed using BBduk from the bbmap suite trimming v38.94 (https://sourceforge.net/projects/bbmap/). STAR Aligner v2.7.3a (Dobin et al. 2013) was used to map the reads to the *Arabidopsis thaliana* genome sequence (Araport11). Reads with identical mapping coordinates and UMI sequences were collapsed to remove PCR duplicates using the dedup command of UMI-tools with the default directional method parameter. Deduplicated reads were counted using HTseq Python framework v0.12.4 (Anders et al. 2015) and using QuantSeq FWD-specified options. The normalization of the raw data and the differential expression analysis were performed on using the DiCoExpress workspace (Lambert et al. 2020; Baudry et al. 2022). We checked that the distribution of the raw p-values followed the quality criterion described by Rigaill et al. (2018). The corresponding R script including the versions of the various R packages is provided in Supplementary Text S1.

### The *ARAGNE* Shiny application a new tool to analyse-omic datasets

Several tools are currently available to analyze, create and visualize gene network in R (i.e. igraph, ggraph, STRINGdb R package, ClusterProfiler, Enrichplot). However, some of the packages focus on statistical analysis or static visualization only. Moreover, the connexion between R and network visualization tool like Cytoscape is mainly manual. Finally, there is a lack of direct integration to obtain reproducible workflow. Starting from this report, we created a user-friendly R shiny application called “Arabidopsis Gene Network Explorer” (*ARAGNE*). This application is a powerful tool designed to analyze and visualize gene networks and perform functional enrichment analysis using GO and KEGG pathways. It integrates with Cytoscape for network visualization and supports correlation analysis, DEG filtering, and enrichment result visualization. Moreover, it allows the production of a comprehensive report to ensure analysis reproducibility. The *ARAGNE* Shiny application is structured as a modular R/Shiny workflow. Users provide (i) normalized gene expression data (gene IDs × samples), (ii) a table of differentially expressed genes (DEG) with at least gene identifiers and log fold-change, and optionally (iii) a list of genes of interest. Data import accepts common tabular formats (e.g. CSV, Excel). The application then allows configurable DEG filtering (log fold-change and adjusted p-value thresholds, choice of up-, down- or both-regulated genes), and correlation analysis between genes (Pearson or Spearman, with user-defined correlation and p-value thresholds and multiple-testing correction via Benjamini–Hochberg or Bonferroni). Built-in visualizations include volcano plots for DEG and interactive correlation summaries. Network construction uses the filtered correlation matrix; nodes can be optionally restricted to genes of interest. Functional enrichment is performed with Gene Ontology (GO) and KEGG pathway analyses via clusterProfiler and org.At.tair.db, and results can be overlaid on the network. Cytoscape integration is achieved through the RCy3 package and the Cytoscape cyREST API, enabling creation and styling of networks, application of layouts, extraction of subnetworks (e.g. neighbourhoods of selected nodes), and export of network images (PNG, PDF, SVG) and session files (.cys) for reproducibility. A report module (R Markdown) produces HTML or PDF reports that record all chosen parameters, summary statistics, correlation and network metrics, and GO/KEGG enrichment results (all html *ARAGNE* reports from the translatome analysis are provided in Supplementary Material S1). *ARAGNE* is freely available, the source code can be obtained from the project repository for local installation and execution. Online documentation, including a user guide and workflow description, is hosted at https://forge.inrae.fr/aragne/ARAGNE. Running *ARAGNE* requires R (version ≥ 4.0), RStudio (recommended), and Cytoscape (version ≥ 3.9) with the cyREST API enabled (default port 1234) for network visualization. Required R packages are listed in the accompanying script Needed_packages.R and include CRAN packages (e.g. shiny, dplyr, igraph, ggplot2, DT, plotly, visNetwork) and Bioconductor packages (RCy3, clusterProfiler, org.At.tair.db, STRINGdb). The application is designed for *Arabidopsis thaliana* gene identifiers; users run it locally in their R environment, ensuring that sensitive data remain on their own machines. No registration or subscription is required to use the tool or to consult the documentation.

### Gene classification by directional pairwise intersection

To identify genes specifically enriched in each of the three translatome datasets (*pLAC17*-associated, *pSWEET17*-associated and *pVND6*-associated), a directional intersection approach was applied to the results of three pairwise differential expression analyses (*pLAC17* vs. *pSWEET17*, *pLAC17* vs. *pVND6*, and *pSWEET17*vs. *pVND6*), performed using edgeR on TMM-normalized log2-counts per million (log2-CPM) values. A gene was retained as a candidate for dataset-specific enrichment if it was called as differentially expressed (false discovery rate, FDR ≤ 0.05) in at least one pairwise comparison. Genes were then classified into two tiers based on the consistency of their differential expression signal. “Clean” genes required bilateral statistical support: a gene was assigned to dataset D*x* only if it was significantly upregulated (FDR ≤ 0.05 and log2 fold-change ≥ 0.5) in both pairwise comparisons involving D*x* — that is, in D*x* vs. D*y* and D*x* vs. D*z* — with concordant directionality. This bilateral criterion ensures that the enrichment signal is reproducible across independent comparisons and is not driven by a single contrast. Genes that were differentially expressed in at least one comparison but did not meet the bilateral criterion — typically because they appeared in only one pairwise comparison or showed inconsistent directionality — were classified as “ambiguous”. Ambiguous genes were assigned to the dataset in which they showed the highest mean log2-CPM expression across biological replicates. This two-tier classification yielded three mutually exclusive groups of dataset-enriched genes (*pLAC17*-, *pSWEET17*-, and *pVND6*-enriched), each comprising a clean subset supported by bilateral evidence and an ambiguous subset assigned by maximum expression. Overlaps between pairwise DEG lists and the composition of each intersection category were visualized using an UpSet plot.

### Visualization of GO enrichment networks in Cytoscape

For each pairwise comparison, the network generated by *ARAGNE* was visualized into Cytoscape. Using Cytoscape’s “Merge” tool, an union network was constructed by integrating the networks derived from both pairwise comparisons of each translatome. This union network was subsequently analyzed using ClueGO against the whole Arabidopsis genome which was used as the reference genes set. The analysis was restricted to Biological Processes (GO:BP) and the network specificity setting to Medium. The GO term fusion option was enabled to group functionally related terms, and only pathways with a p-value below 0.05 were retained for visualization. For statistical testing, a two-sided hypergeometric test was employed, with p-value correction performed using the Bonferroni step-down method. The resulting network was visualized using an “organic” layout, which optimizes edge length to minimize overlaps, enhance clarity, and facilitate the visual distinction of functional clusters. In the final GO enrichment network, node size is proportional to the number of genes associated with each GO term, while edge thickness reflects the Kappa score, representing the degree of similarity between terms based on shared gene overlap. In addition, GO enrichment networks were performed using the list of genes enriched in each translatome and analyzed using ClueGO with the same settings than those described above.

### Plant materials and growth conditions

The following Arabidopsis lines have been used in this work: Columbia-0 accession (used as wild-type plants and hereafter referred as WT), the *sweet2-1*, *sweet16-4*, *sweet17-1* single mutant and the *sweet16-4sweet17-1* double mutant, previously characterized (Chen et al. 2015; Aubry et al. 2022) (hereafter referred as *swt2*, *swt16*, *swt17*, and *swt16;17* respectively), the *sweet2-1sweet16-4*, the *sweet2-1sweet17-1* double mutants and the *sweet2-1sweet16-4sweet17-1* triple mutant (hereafter referred as *swt2;16, swt2;17* and *swt2;16;17*, respectively) were obtained by crossing. The complemented line was obtained by crossing the *swt2* single mutant lines with the corresponding translational fusions lines pSWEET2:SWEET2-GUS (Chen et al. 2015). Homozygous plants were genotyped using gene-specific primers in combinations with a specific primer for the left border to the T-DNA insertion (Supplementary Table S11). In addition, the expression of *SWEET2*, *SWEET16* and *SWEET17* was verified by qPCR in the *swt2;16;17* triple mutant line (Supplementary Fig. S10). As expected, a significantly low expression of the three genes was measured in the triple mutant line compared to wild-type plants (Supplementary Fig. S10). Seeds were stratified at 4°C for 72 hours in 0.1 % agar solution to synchronize germination. Plants were grown in a growth chamber in long-day conditions as described in Aubry et al. (2022). For all experiments, the main inflorescence stems (after the removal of lateral inflorescence stems, flowers and siliques) were harvested from 45 DAS plants.

### Growth parameters

Projected rosette area (PRA) was measured from pictures taken at 35 and 45 DAS and the use of FIJI software (Schindelin et al. 2012). The stem height and diameter were measured at 45 DAS using a ruler and a digital caliper, respectively.

### Morphometric analysis of the stem tissues

A 1- to 2-cm segment taken at the bottom part of the main inflorescence stem was harvested. The stem segments were embedded in 8% (w/v) agarose solution and sectioned using a VT100S vibratome (Leica, https://www.leica-microsystems.com). Some of the cross-sections were used for FT-IR analysis (for details see dedicated paragraph below) and the other were stained with a FASGA staining solution (Tolivia and Tolivia 1987). Stained inflorescence stem cross-sections were imaged under an Axio Zoom V16 microscope equipped with a Plan-Neofluar Z 2.3/0.57 FWD 10.6 objective (Zeiss). Since there is a heterogeneity in vascular bundle shape within one inflorescence stem section (Hoffmann et al. 2022), only the shape of the M-type vascular bundles were measured. For each section, several parameters were measured: stem diameter, the phloem area of each M-type vascular bundle, the xylem area of each M-type vascular bundle and the interfascicular fibers thickness. In addition, the xylem tissue shape was assessed by measuring the tangential and radial length of the vascular bundle as well as the tangential:radial length ratio. The tangential length is used as a proxy for difference in xylem size while the radial length was measured as a proxy for difference in xylem cell number according to Wang et al. (2019). All parameters were measured using the FIJI software (Schindelin et al. 2012).

### Targeted gene expression analysis

RNAs were prepared from the main inflorescence stem of four individual 45 DAS plants. Tissue collection, RNA extraction, DNase treatment, and cDNA synthesis were performed as described in Aubry et al. (2022). RT-qPCR analyses were conducted on four independent biological replicates, each with three technical replicates, using gene-specific primers from the literature (Supplementary Table S11). Reactions were carried out in 10-µL volumes on a Bio-Rad CFX96 Real-Time PCR system using Takyon ROX SYBR MasterMix, with cycling and melting-curve conditions as described in Aubry et al. (2022). Cq values were obtained with Bio-Rad CFX Manager 3.0; technical replicates differing by more than 0.5 Cq were excluded. Among four candidate reference genes (*APT1*, *TIP41*, *EF1a*, and *UBQ5*), *APT1* was identified as the most stable using NormFinder and used for normalization. Relative transcript levels were calculated using the ΔCt method, accounting for primer efficiencies.

### Histochemical GUS staining

The translational fusion between *GUS* and the C-terminus of *SWEET2* genomic sequence under the control of its respective native promoter (2546 bp) in a wild-type background was kindly provided by Dr. Woei-Jiun Guo (National Cheng Kung University, Tainan, Taiwan) (Chen et al. 2015). The expression pattern of both SWEET2 and the transcriptional fusions *GENE_pro_:HF-RPL18-GUS* were assessed at the bottom part of the main inflorescence stem of 45 DAS plants grown in growth chamber conditions. The histochemical GUS staining was performed according to Sorin et al. (2005). Inflorescence stem subjected to GUS staining were then embedded in 8% (w/v) agarose and sectioned using a Leica VT100S vibratome (Leica, htpps://www.leica-microsystems.com/). Sections were counterstained for lignin by phloroglucinol staining (Pradhan Mitra and Loqué 2014). Pictures were taken using a Leitz Diaplan microscope equipped with an AxioCam MRc camera and the ZEN (blue edition) software package (Zeiss, https://www.zeiss.com/).

### Soluble sugars and starch quantification

The main inflorescence stems and the rosette leaves of the wild-type, and the *swt2*, *swt16*, *swt17*, *swt2st16*, *swt2swt17*, *swt16swt17* and *swt2swt16swt17* mutants, were harvested in the middle of the day (8 h after the beginning of the light period), frozen in liquid nitrogen, and ground with a mortar and a pestle. Samples were then freeze-dried using a freeze dryer for 24 hours. Soluble sugars and starch were extracted from 15-19 mg of dry powder as described in Aubry et al. (2022, 2024).

### FTIR analysis of the secondary cell wall

On the previously cut stem cross-sections (see paragraph “Morphometric analysis of the stem tissues”), the composition of the secondary cell wall of the xylem tissues and the interfascicular fibers were determined by Fourier Infra-Red spectroscopy using an FT-IR Nicolet iN (Thermo Fisher Scientific, https://www.thermofisher.com). Spectral acquisition were done in reflection mode on a 30 µm x 30 µm acquisition targeting the xylem tissue (including both xylem vessels and fibers) or the interfascicular fibers, as described in Le Hir et al. (2015). Between 14 and 25 spectra sweeping the xylem tissue or the interfascicular fiber homogeneously were acquired per plant for each genotype on one stem cross-section. Between nine and twelve individual inflorescence stems coming from two independent cultures were analyzed for each genotype. After sorting the spectra and correcting the baseline, the spectra were area-normalized and the different genotypes were compared as described in Le Hir et al. (2015). The absorbance values (maximum height) of the major cellulose, lignin and hemicellulose bands in the fingerprint region (1,800–800 cm^−1^) were collected on baseline-corrected and normalized spectra using the ‘hyperSpec’ R package (Beleites and Sergo 2020).

### Statistical analysis

To test for differences between the condition/genotypes, one-way ANOVA were performed. Tukey (HSD) post-test was used as it allows for a rigorous control of type I error (false positive) in multiple comparisons between all pairs of means, thus ensuring statistical robustness. Alternatively, a Duncan post-test was used to maximize the sensitivity of detecting differences between groups. Finally, a student *t*-test performed on Microsoft Excel software was used to compare the different lines to the wild-type plants. Principal component analysis was performed using FactoMineR package of R (Le et al. 2008). All the statistical analysis were done using R (version 4.4.2) and Rstudio (version 2025.09.2+418).

### Accession numbers

Arabidopsis Genome Initiative locus identified for each gene are as follows: *SWEET2* (At3g14770), *SWEET16* (At3g16690), *SWEET17* (At5g60020), *APT1* (At1g27450), *TIP41* (At4g34270), *EF1*α (At5g60390), *UBQ5* (At3g62250).

## Supporting information

Supplementary Data Set1

Supplementary Data Set2

Supplementary Data Set3

Supplementary Data Set4

Supplementary TablesS1-S11

## ACKNOWLEDGMENTS

This work has benefited from the support of IJPB’s Plant Observatory platforms PO-Plants, PO-Cyto and PO-Chem. This work has benefited from a French State grant (Saclay Plant Sciences, reference n° ANR-17-EUR-0007, EUR SPS-GSR) managed by the French National Research Agency under an Investments for the Future program integrated into France 2030 (reference n° ANR-11-IDEX-0003-02).

## AUTHOR CONTRIBUTIONS

R.L.H. conceptualized the research. B.H., F.V., A.L.A, D.M., S.L. and M.Y. performed the experiments and the data analysis. The data visualization was performed by R.L.H. The original draft writing was performed by R.L.H. and the final review and editing was performed by all authors. All authors agreed with the final version.

## DATA AVAILABILITY

The RNA-seq raw data reported in this work has been deposited in the Gene Expression Omnibus (GEO) website (https://www.ncbi.nlm.nih.gov/geo/) under the accession number GSE322629.

## SUPPLEMENTARY DATA

**Supplementary Figure S1.** PCA plot of all datasets derived from translatome analysis of three stem tissues.

**Supplementary Figure S2.** Differentially expressed gene overlaps across pairwise translatome comparisons.

**Supplementary Figure S3.** GO term enrichment analysis of upregulated genes in each pairwise translatome comparisons.

**Supplementary Figure S4.** Heatmap of enriched genes across three translatome datasets.

**Supplementary Figure S5.** Expression of cambium-related genes in the different translatomes.

**Supplementary Figure S6.** GO enrichment network of enriched genes in *pSWEET17*-associated translatome.

**Supplementary Figure S7.** *SWEET2*-centered subnetworks identified from translatome comparisons.

**Supplementary Figure S8.** Phenotypic characterization of the *sweet* mutant series.

**Supplementary Figure S9.** Cell wall composition of xylem secondary cell wall and interfascicular fibers.

**Supplementary Figure S10.** The expression of *SWEET2*, *SWEET16* and *SWEET17* are significantly downregulated in the *swt2;16;17* triple mutant.

**Supplementary Table S1.** Percentage of translated genes in the different xylem cell types and interfascicular fibers.

**Supplementary Table S2.** GO enrichment analysis of enriched genes in *pVND6*-associated translatome compared to *pLAC17*- and *pSWEET17*-associated translatomes.

**Supplementary Table S3.** GO enrichment analysis of enriched genes in *pSWEET17*-associated translatome compared to *pVND6*- and *pLAC17*-associated translatomes.

**Supplementary Table S4.** Expression of transcripts specifically enriched in the *pSWEET17*-associated translatome in single-cell RNA-seq data from Arabidopsis roots undergoing secondary growth.

**Supplementary Table S5.** GO enrichment analysis of genes enriched in *pSWEET17*-associated translatome and found with higher average expression in xylem parenchyma cells using single-cell transcriptomic data.

**Supplementary Table S6.** GO enrichment analysis of genes enriched in *pSWEET17*-associated translatome and found with higher average expression in xylem vessels using single-cell transcriptomic data.

**Supplementary Table S7.** GO enrichment analysis of commonly enriched genes in *pLAC17*-associated translatome compared to *pVND6*- and *pSWEET17*-associated translatomes.

**Supplementary Table S8.** List of genes belonging to sugar metabolism, signaling and transport queried in the different xylem translatomes.

**Supplementary Table S9.** Expression of target genes related to sugar metabolism, transport and signaling in a published transcriptomic dataset.

**Supplementary Table S10.** Genes of the *SWEET2*-centered network and their associated expression and network topology parameters.

**Supplementary Table S11.** Primers used for characterizing mutant lines.

**Supplementary Data Set 1.** RNA-seq raw and normalized counts.

**Supplementary Data Set 2.** Differentially expressed gene in pairwise comparisons of the three translatomes.

**Supplementary Data Set 3.** Lists of differentially expressed gene overlaps across pairwise translatome comparisons.

**Supplementary Data Set 4.** Lists of genes enriched in each translatome after directional intersection approach

**Supplementary Text S1.** R script and the versions of the different R packages used for the DiCoExpress analysis.

**Supplementary Material S1**. Zip file including .html *ARAGNE* analysis reports.

## Supplementary Figures

**Supplementary Figure S1.**
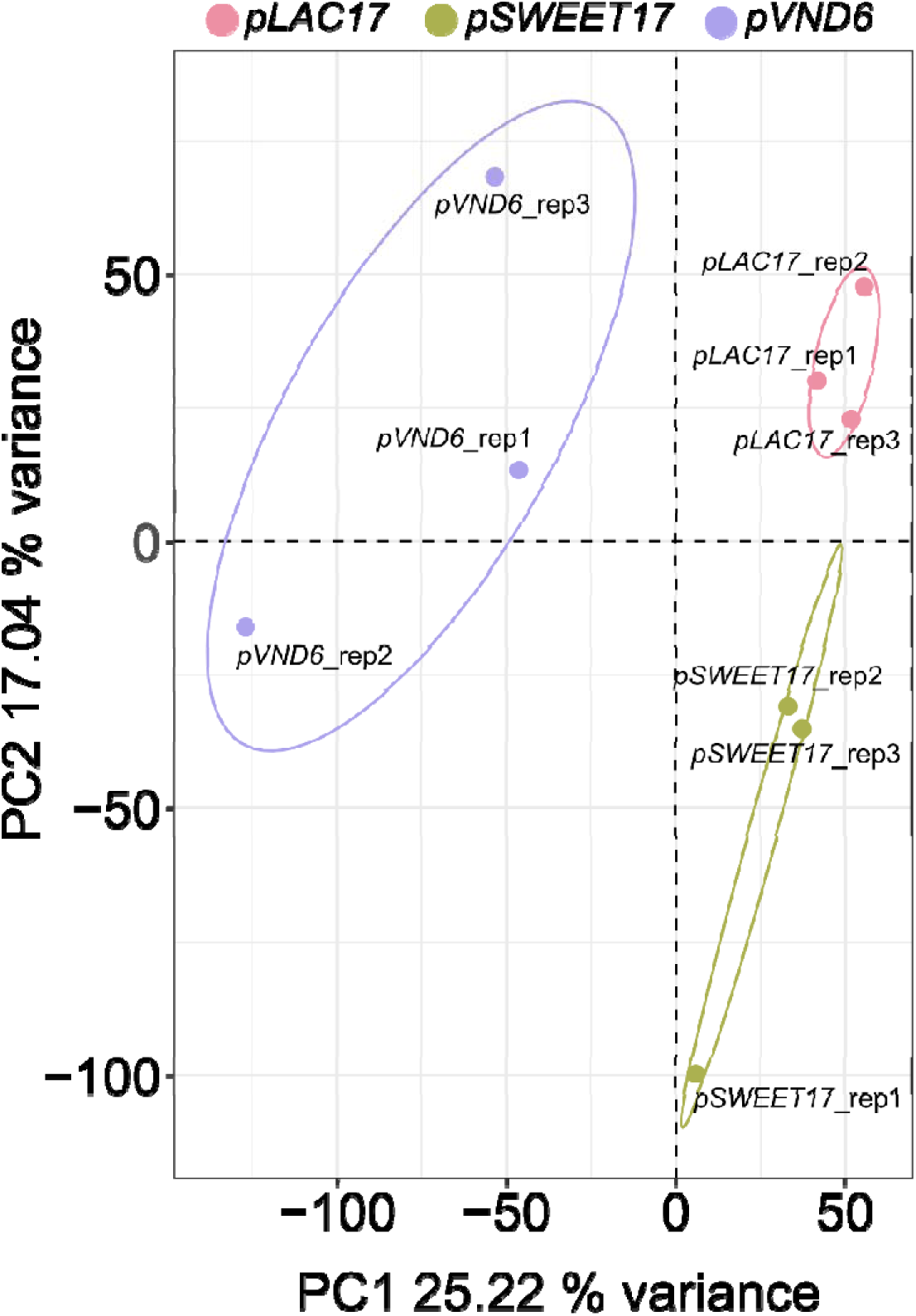
PCA plot of all datasets derived from translatome analysis of three stem tissues. PCA of log_2_-transformed normalized read counts of each RNA-seq datasets.

**Supplementary Figure S2.**
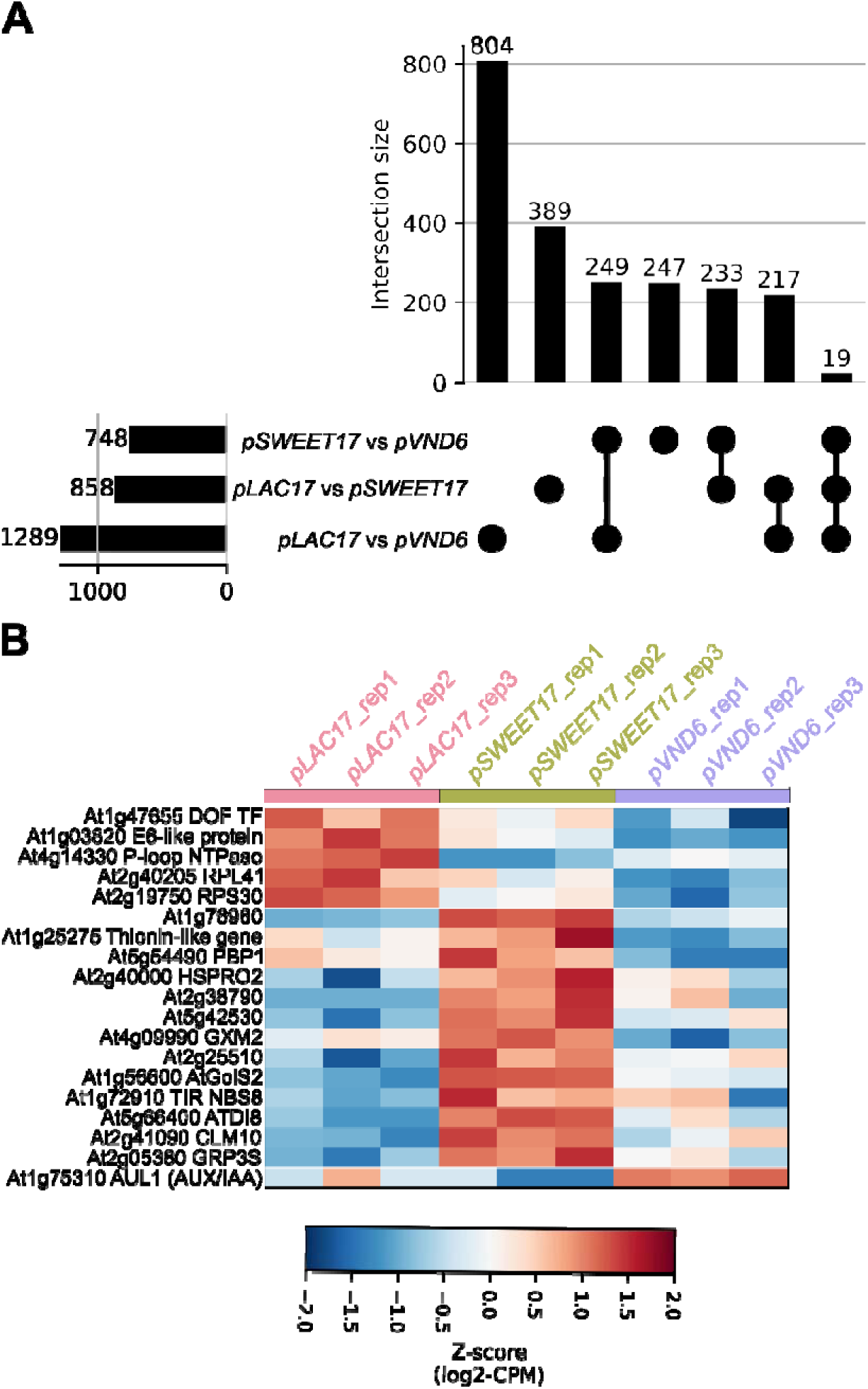
Differentially expressed gene overlaps across pairwise translatome comparisons. (A) Upset plot showing the number of genes overlapping across the pairwise comparisons. The horizontal bars (left) show the total number of DEGs identified in each pairwise comparison (*pLAC17* vs *pVND6*: 1289; *pLAC17* vs *pSWEET17*: 858; *pSWEET17* vs *pVND6*: 748; FDR ≤ 0.05, |log2FC| ≥ 0.5). The vertical bars (top) show the size of each intersection category, with filled circles and connecting lines indicating which comparisons contribute to each intersection. (B) Heatmap showing the normalized expression of the genes differentially expressed in all three pairwise comparisons simultaneously. Color intensity represents the Z-score of log2-CPM values across the nine samples (three biological replicates per dataset). Genes are grouped by their assigned dataset and ordered by hierarchical clustering (Ward linkage, Euclidean distance) within each group. The list of differentially expressed genes of each pairwise comparison and intersections can be found on Supplementary Data Set 2 and Data Set 3.

**Supplementary Figure S3.**
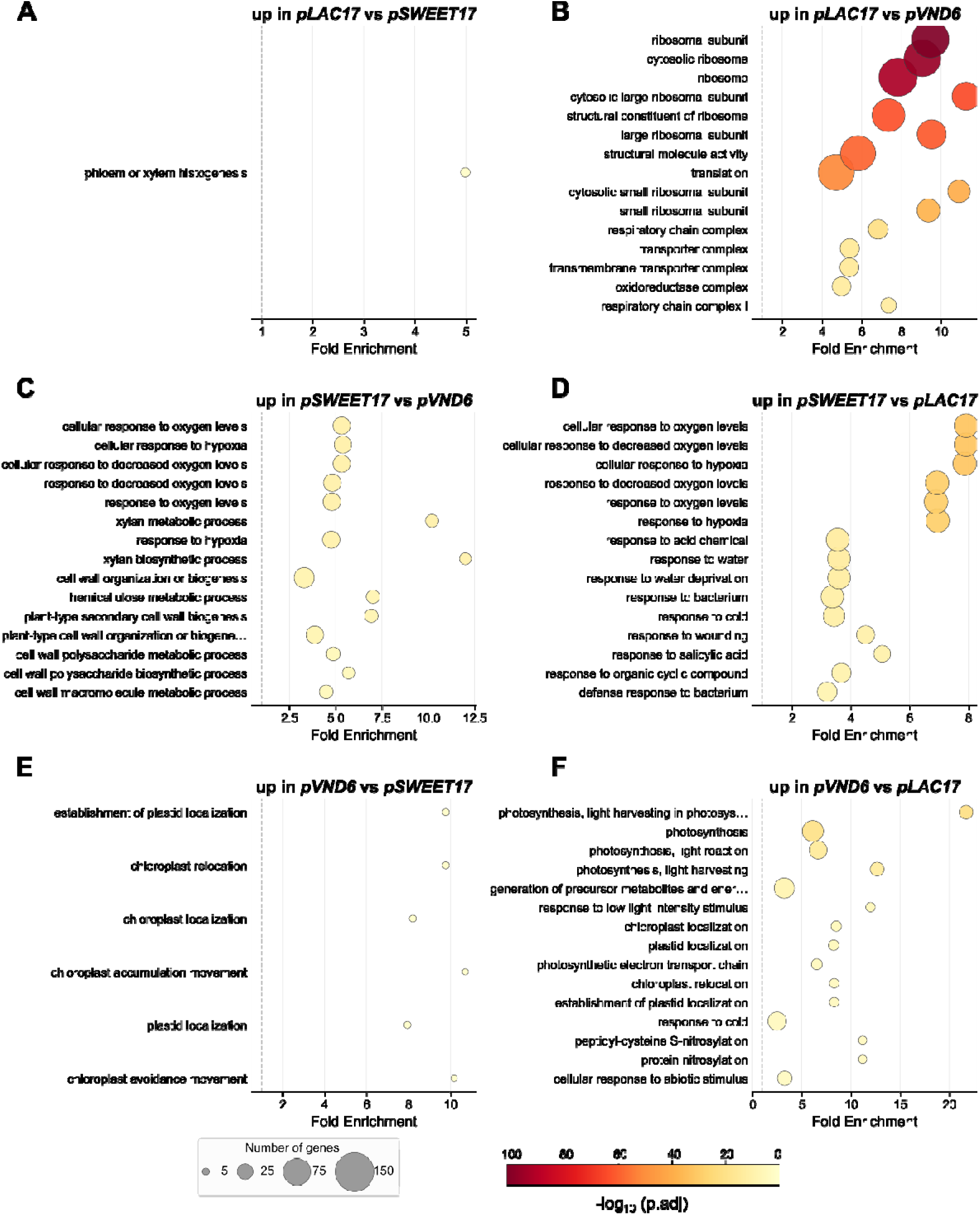
GO term enrichment analysis of upregulated genes in each pairwise translatome comparisons. Bubble plots showing the top 15 over-represented Gene Ontology (GO) Biological Process terms for genes upregulated in each pairwise comparison between the three promoter-driven associated translatomes (*pLAC17*, *pSWEET17*, *pVND6*). Each bubble represents one GO term. The position along the x-axis indicates the fold enrichment of the term relative to the whole genome background. Bubble size is proportional to the number of genes annotated to that GO term in the input gene list (gene count). Bubble color reflects statistical significance, expressed as −log□□(adjusted p-value), on a yellow-to-dark-red scale; darker red indicates greater significance. Adjusted p-values were computed using the Benjamini–Hochberg method. Terms are ordered from bottom to top by decreasing adjusted p-value (most significant at top). Terms with adjusted p-values comprised between 0.05 and 0.10 were retained; panels with few bubbles reflect comparisons with limited enrichment signal. The *ARAGNE* .html files for this analysis can be found in Supplementary Material S1 and the lists of differentially expressed genes for each pairwise comparison in Supplementary Data Set 2.

**Supplementary Figure S4.**
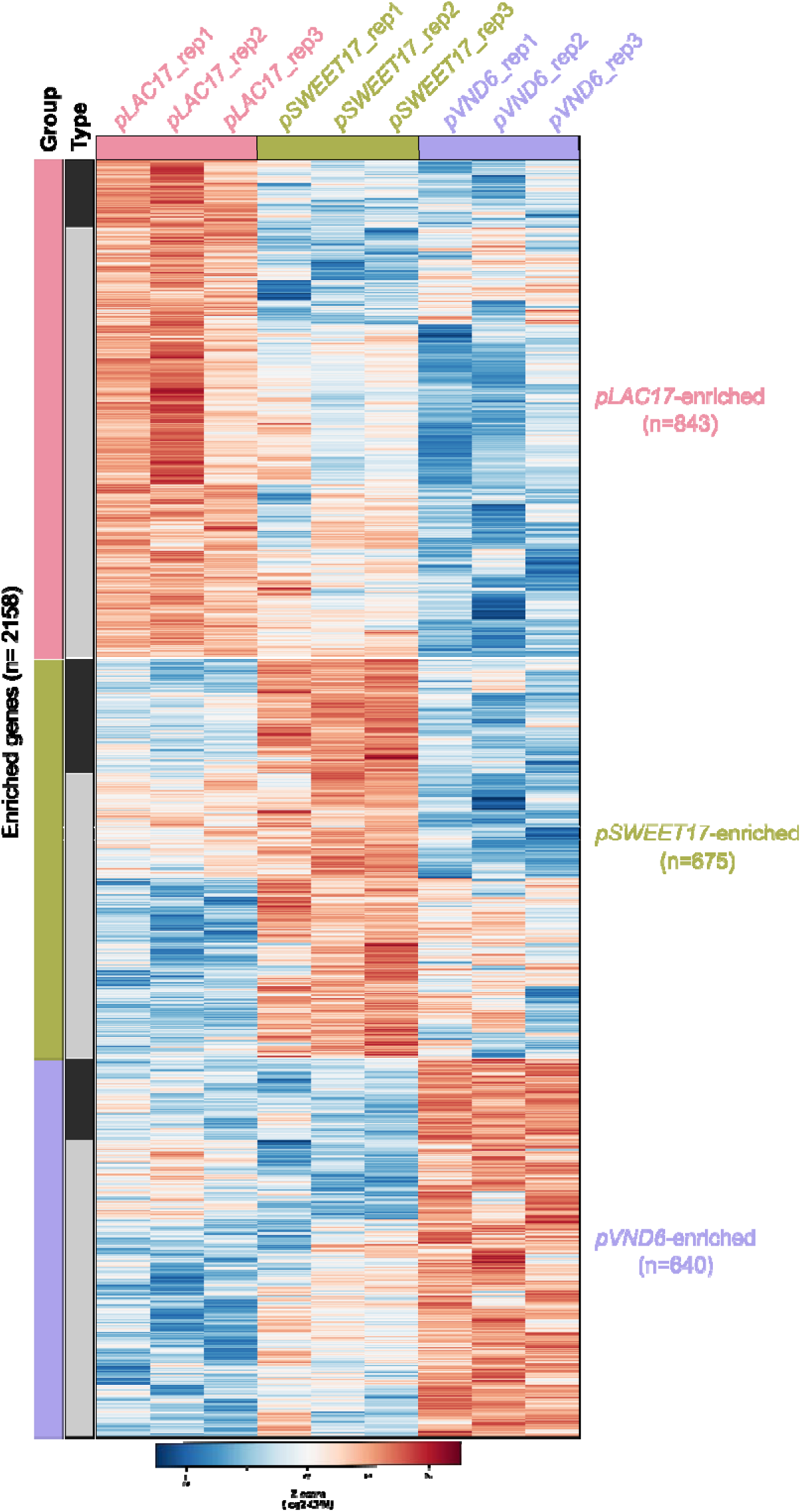
Heatmap of enriched genes across the three translatome datasets. Heatmap showing the normalized expression of 2,158 differentially expressed genes (DEGs) enriched in *pLAC17*-associated translatome (n = 843 of which 116 were common in both relevant comparisons), *pSWEET17*-associated translatome (n = 675 of which 195 were common in both relevant comparisons), or *pVND6*-associated translatome 3 (n = 640 of which 138 were common in both relevant comparisons). Color intensity represents the Z-score of log2-CPM values across the nine samples (three biological replicates per dataset). Genes are grouped by their assigned dataset and ordered by hierarchical clustering (Ward linkage, Euclidean distance) within each group. The left annotation strips indicate the assigned dataset (color) and classification type (dark: genes identified by pairwise intersection logic in both relevant comparisons; light grey: genes assigned to the dataset with the highest mean expression). The list of genes enriched in each translatome can found on Supplementary Data Set 4.

**Supplementary Figure S5.**
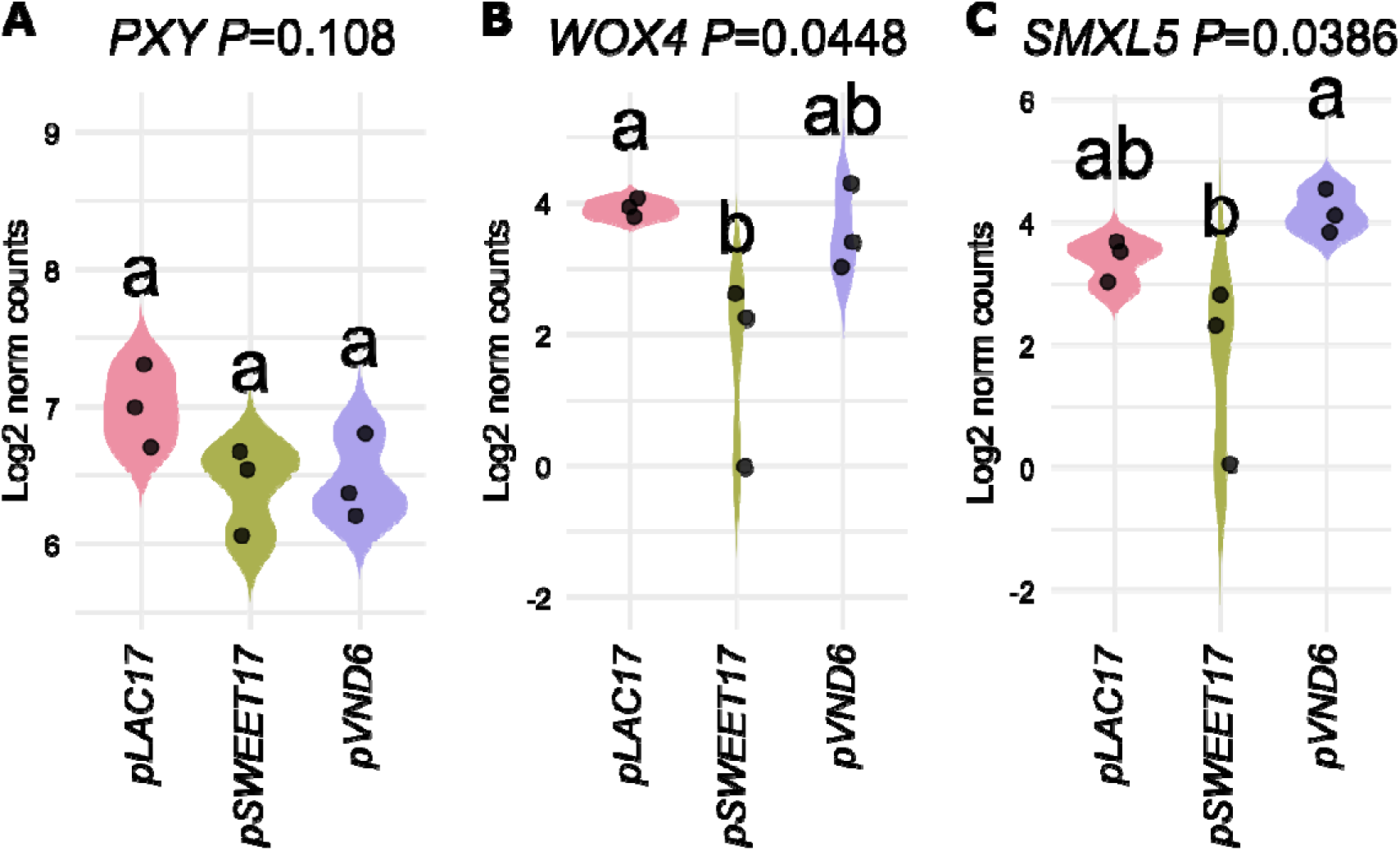
Expression of cambium-related genes in the different translatomes. **(A-C)** Violin plots showing log_2_-transformed normalized gene reads counts of the *PXY* (A), *WOX4* (B) and *SMXL5* (C) among the different translatomes. Each black dot corresponds to a biological replicate (n = 3). A one-way ANOVA combined with Tukey’s comparison post-test has been made to compare the different translatomes. The different letters indicate significant differences. The *P*-value for the comparison between the different translatomes is indicated on the graph.

**Supplementary Figure S6.**
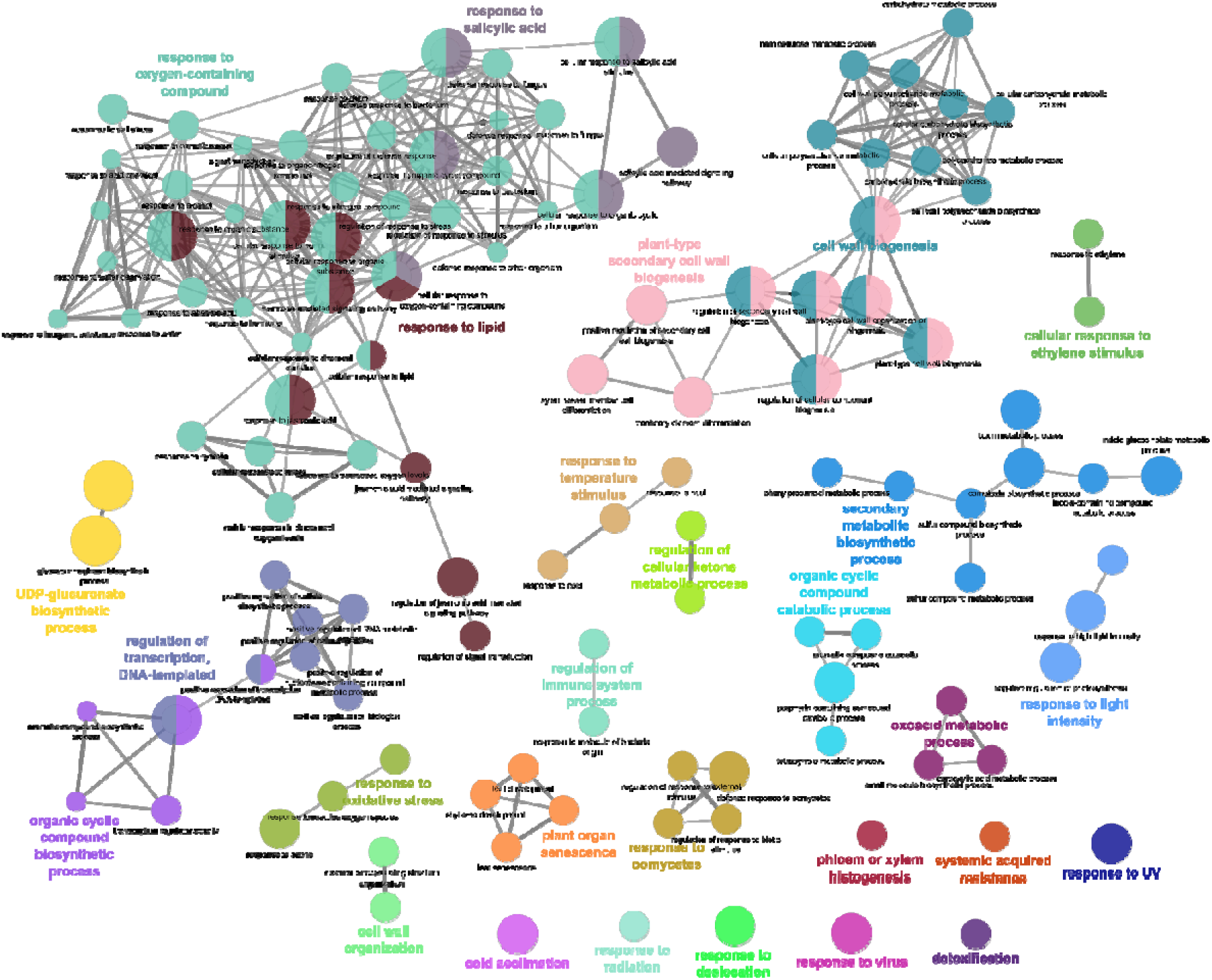
GO enrichment network of enriched genes in *pSWEET17*-associated translatome. Gene Ontology (GO) enrichment analysis was performed on the set of genes identified as enriched in *pSWEET17*-associated translatome in comparison with *pLAC17*- and *pSWEET17*-associated genes sets. The list of genes is provided in Supplemental Data Set 4. Enrichment analysis was conducted using ClueGO and visualized in Cytoscape, as detailed in the *Materials and Methods*. Node size reflects the number of genes associated with each GO term in the TAIR11 background annotation and edges thickness is related to the KAPPA score. Only GO terms related to Biological Processes (BP) and for which a p-value lower than 0.05 are shown. The full list of enriched GO terms is available in Supplementary Table S3.

**Supplementary Figure S7.**
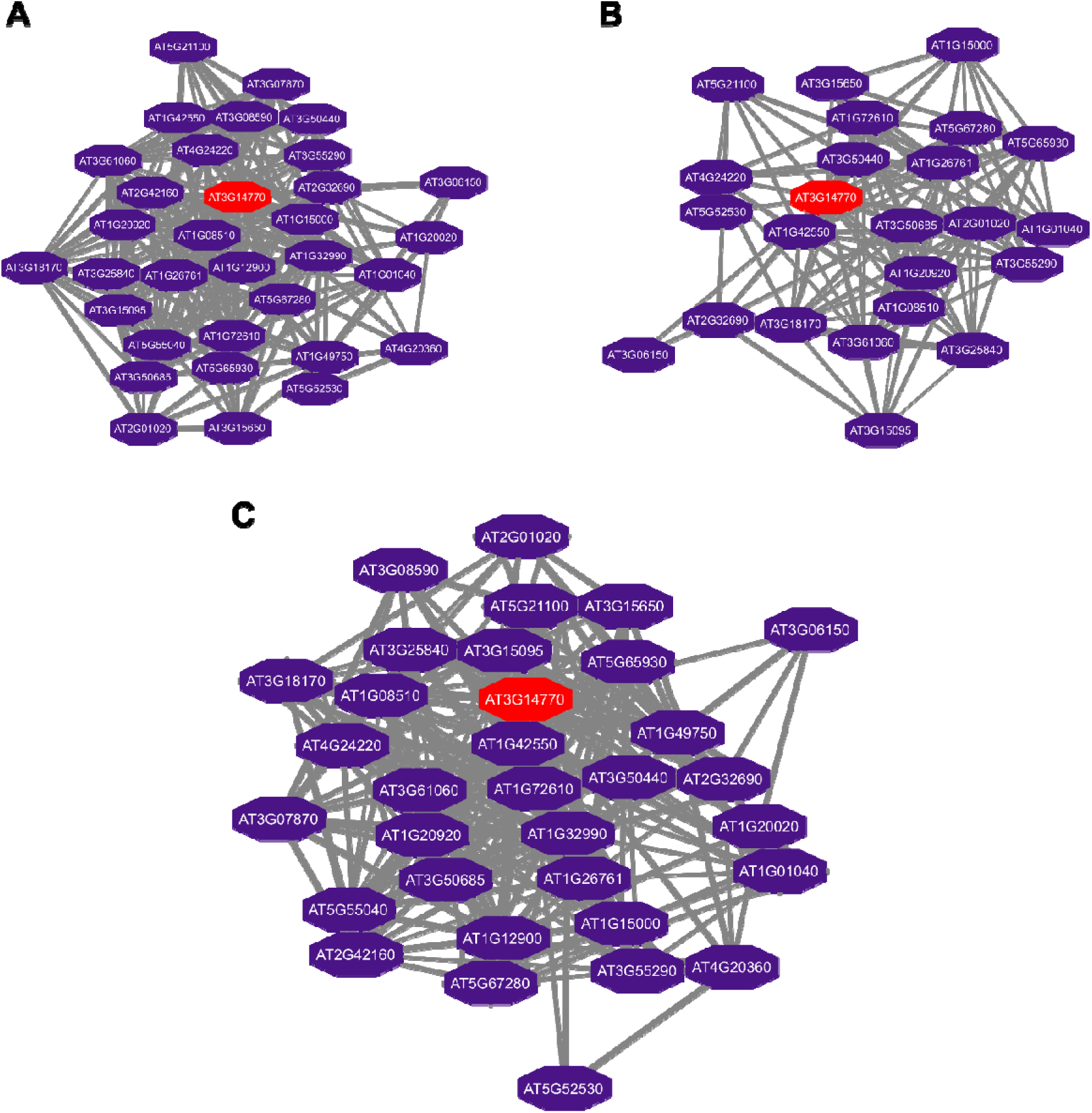
*SWEET2*-centered subnetworks identified from translatome comparisons. (A) subnetwork extracted following differential analysis between. *LAC17*-and *VND6*-associated translatomes. (B) subnetwork extracted following differential analysis between *SWEET17*- and *VND6*-associated translatomes. (C) Merged network produces from the Union of both subnetworks (A and B).

**Supplementary Figure S8.**
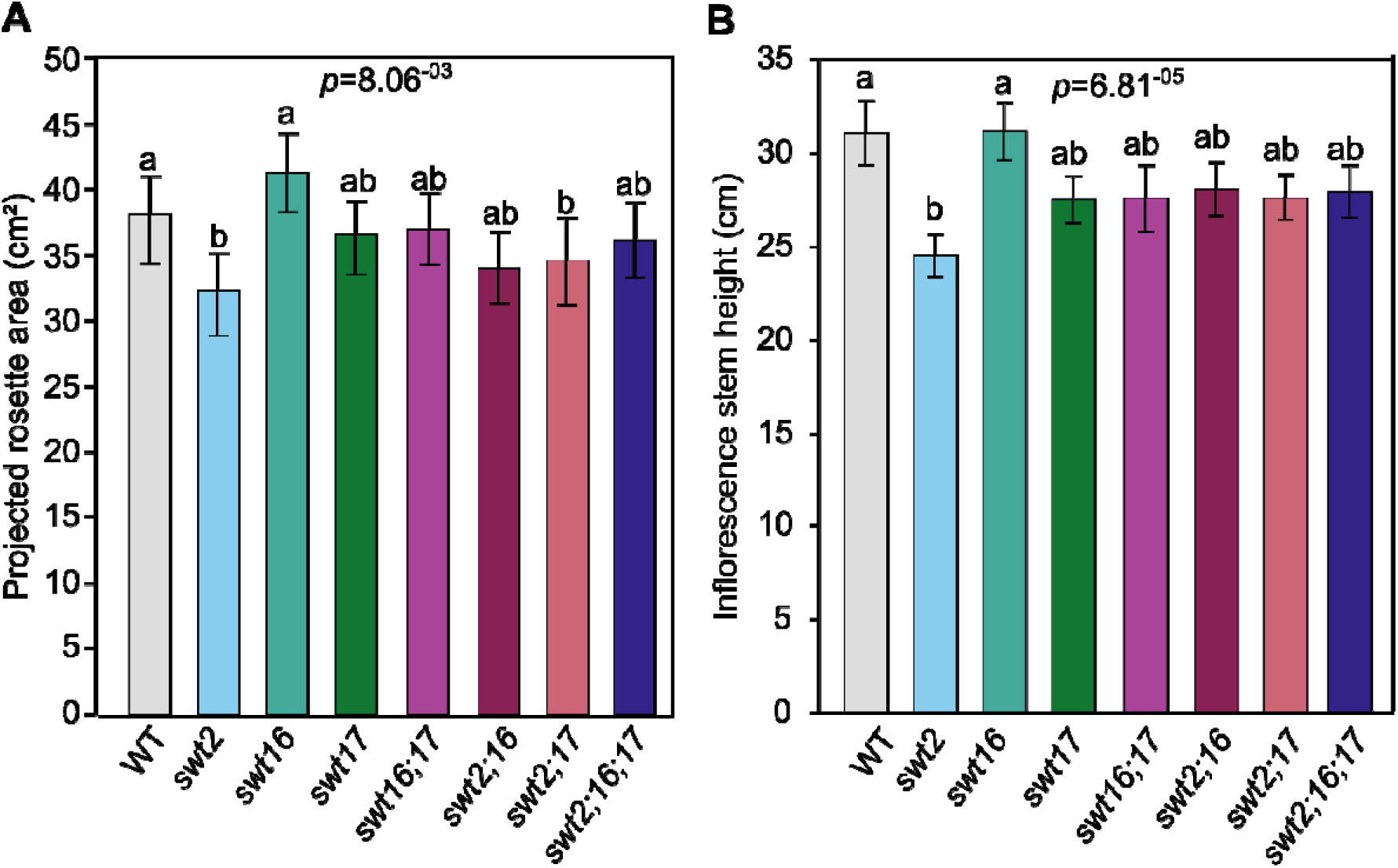
Phenotypic characterization of the *sweet* mutant series. Barplots showing projected rosette area at 35 DAS (A) and inflorescence stem height at 45 DAS (B). Least-square means from two independent experiments ± SE are shown (n ≥ 10 for each genotype). A one-way ANOVA combined with Tukey’s comparison post-test has been made to compare the different genotypes. The different letters indicate significant differences. The *P*-values for the comparison between the different genotypes are indicated on each graph.

**Supplementary Figure S9.**
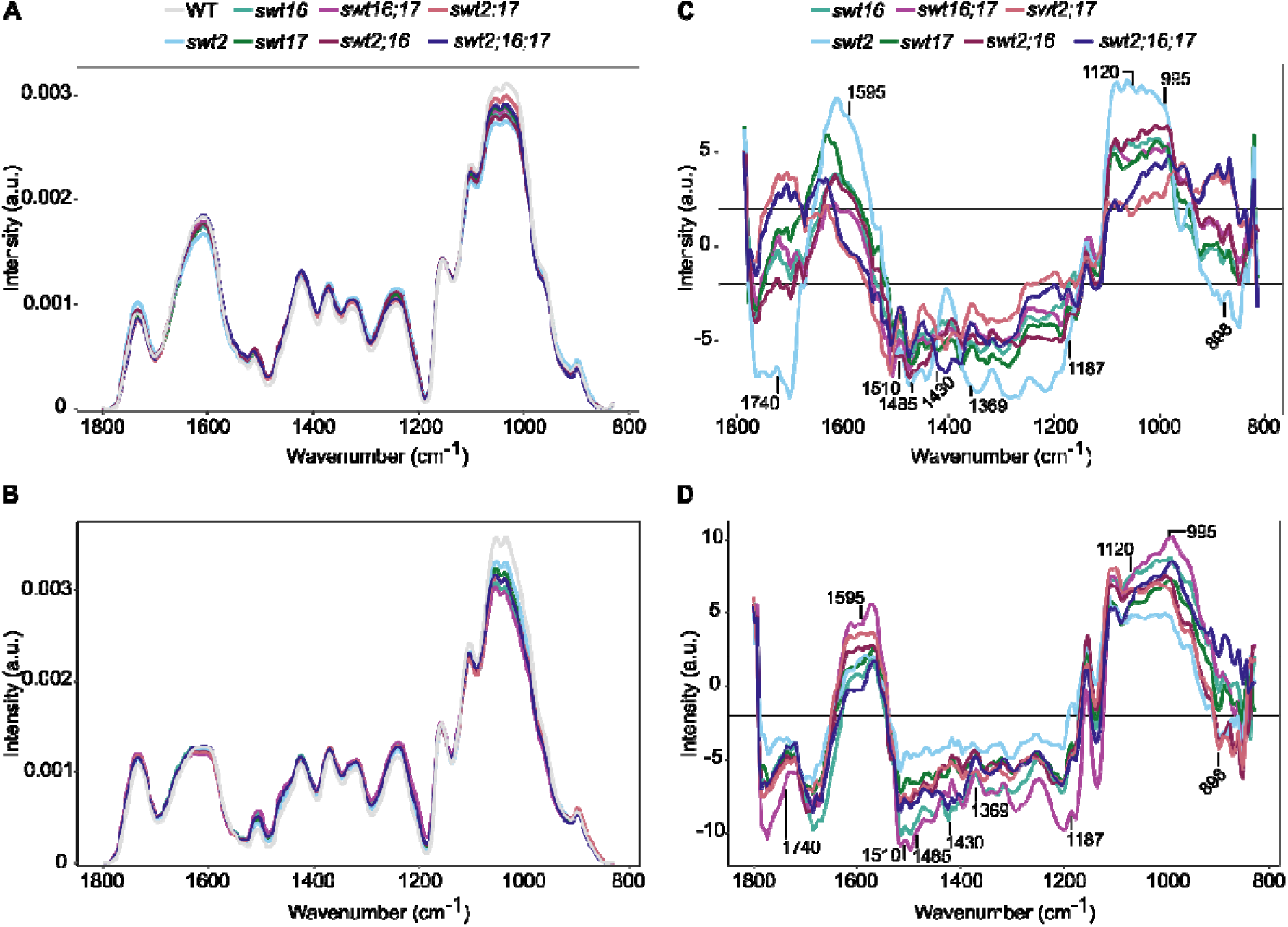
Cell wall composition of xylem secondary cell wall and interfascicular fibers. FTIR spectra were acquired on xylem tissue and on interfascicular fibers from sections of the basal part of the inflorescence stem. All spectra were baseline-corrected and area-normalized in the range 1800–800 cm^−1^. A, Average FTIR spectra from xylem tissue were generated from 272, 186, 127, 182, 139, 133, 168, and 205 spectra for wild-type, *swt2*, *swt16*, *swt17*, *swt16;17*, *swt2;16*, *swt2;17* and *swt2;16;17* plants, respectively, obtained using 9-12 independent plants for each genotype grown in two independent cultures. B, Average FTIR spectra from interfascicular fibers were generated from 253, 249, 143, 188, 146, 168, 167, and 208 spectra for wild-type, *swt2*, *swt16*, *swt17*, *swt16;17*, *swt2;16*, *swt2;17* and *swt2;16;17* plants, respectively, obtained using 9-12 independent plants for each genotype grown in two independent cultures. C-D, Comparison of FTIR spectra obtained from xylem cells (C) and interfascicular fibers (D) of the *swt2*, *swt16*, *swt17*, *swt16;17*, *swt2;16*, *swt2;17* and *swt2;16;17* mutants. A Student’s t test was performed to compare the absorbances for the wild-type, single, double and triple mutants. The results were plotted against the corresponding wavenumbers. *t*-Values (vertical axis) between –2 and + 2 correspond to nonsignificant differences (*P* < 0.05) between the genotypes tested (9 > n < 12). *t*-Values above + 2 or below –2 correspond to, respectively, significantly weaker or stronger absorbances in the mutant spectra relative to the wild-type.

**Supplementary Figure S10.**
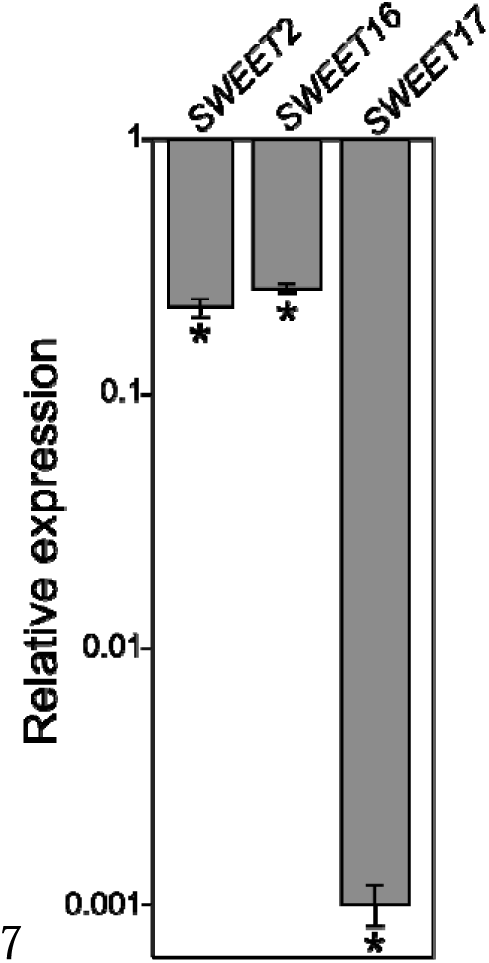
The expression of *SWEET2*, *SWEET16* and *SWEET17* are significantly downregulated in the *swt2;16;17* triple mutant. mRNAs were extracted from wild-type plants grown in long-day conditions at 22°C for 45 days. The mRNA contents are expressed relative to those of *APT1* and normalized to the value of wild-type plants, which was arbitrarily set to 1. Values represent means ± SD from three biological replicates. Asterisks indicate a significant difference (* *P* < 0.05) from wild type according to Student’s *t* test.

