## Supplementary Material S1 for "Translational profiling uncovers a tonoplast sugar transporter essential for vascular system development and cell wall composition": ARAGNE_report_up_pSWEET17_comp_pLAC17.html

Network Analysis Report


### Network Analysis Report

###### Generated by Shiny Application

#### 2026-04-14

- 1 Introduction
- 2 Analysis Parameters
- 3 Analysis Summary
  - 3.1 Differentially Expressed
    Genes
  - 3.2 Network Metrics
- 4 Correlation Table
- 5
  Network Metrics Tabel
- 6 Visualizations
- 7 Conclusion

### 1 Introduction

This report summarizes the results of a network analysis and
functional enrichment study, with a focus on differentially expressed
genes and correlation-based network metrics.

---

### 2 Analysis Parameters

Here are the key parameters used in the analysis:

- **LogFC Threshold:** 0
- **P-value Threshold for Differential Expression
  Analysis:** 0.05
- **P-value Threshold for Correlation Analysis:**
  0.05
- **Correlation Method:** pearson
- **P-value Adjustment Method:** BH
- **Regulation Type:** down

---

### 3 Analysis Summary

#### 3.1 Differentially Expressed Genes

- **Total number of differentially expressed genes**:
  858
- **Upregulated Genes**: 314
- **Downregulated Genes**: 544

#### 3.2 Network Metrics

- **Total Nodes**: 535
- **Total Edges**: 5499
- **Average Betweenness**: 613.35
- **Max Correlation**: 1
- **Min Correlation**: 0.9
- **Overlap with Genes of Interest**: 1
- **Genes of Interest**: AT1G08930

### 4 Correlation Table

Below is the correlation table showing the correlations among
differentially expressed genes.

### 5 Network Metrics Tabel

The chosen network for visualization in Cytoscape is:
**Main\_Network-upSWT17-vs-pLAC17**.

A summary of the network metrics is provided in the table below.

### 6 Visualizations

```
## The chosen network for GO Enrichment Analysis is: Main_Network-upSWT17-vs-pLAC17
```

```
## The chosen network for GO Enrichment Analysis is: Main_Network-upSWT17-vs-pLAC17
```

### 7 Conclusion

This analysis provides insights into the network structure and gene
relationships within the study. Based on the network metrics and
correlation analysis, further steps could include deeper functional
enrichment analysis or visualization in Cytoscape.
